# Topographical archetypes of somatic mutagenesis in cancer

**DOI:** 10.64898/2026.04.18.719374

**Authors:** Allen W. Lynch, Sangmi Sandra Lee, Jan P. Hummel, Benedikt Geiger, Michael S. Lawrence, Hu Jin, Doga C. Gulhan, Peter J. Park

## Abstract

The genome of every cancer cell carries a record of the mutational processes that have acted throughout its history. Mutational signature analysis, which infers the activity of mutagenic processes from their characteristic base-change patterns, has become an indispensable tool for interpreting somatic mutations. However, this framework captures only which types of mutations a process generates and not where in the genome they occur — a distribution influenced by replication timing, chromatin organization, transcription, DNA secondary structure, and other genomic features. Here, we present a generative probabilistic framework (MuTopia) that jointly infers mutational spectra and their genome-wide topography as nonlinear functions of genomic and epigenomic state. Applied to whole-genome sequencing data from 15 tumor types, MuTopia reveals that mutational processes fall into eight conserved topographic archetypes, or topotypes, shaped primarily by replication timing and chromatin state. Diverse mutational processes converge upon this limited repertoire, indicating that the genomic distribution of mutagenesis is constrained less by the source of damage than by how that damage is processed. Individual mutational processes exhibit state-dependent variation in their genomic distributions: the same signature can adopt distinct topotypes depending on repair proficiency and replication stress. For instance, SBS8 shifts from a canonical late-replicating profile in homologous recombination-proficient tumors to an early-replicating, stress-associated topotype in HR-deficient tumors, and replication stress similarly reshapes the genomic distribution of APOBEC editing. Topotypes, therefore, provide a classification of mutagenesis distinct from spectral signatures, capturing aspects of tumor biology that spectra alone cannot resolve.

## Introduction

Mutational processes encode the evolutionary history of somatic cells into their genomes. As cells grow, divide, and age, they accumulate genetic insults — DNA damage, erroneous repair or replication, and aberrant enzymatic activity — that leave characteristic base changes in the genome. From these patterns, one can decipher the etiology and molecular characteristics of cancer, from prior mutagen exposure [1–4] to DNA repair deficiencies [5–8], with implications for prevention and treatment [9].

The mutational signatures framework leverages this notion: a “signature” describes the rate at which a process generates mutations across a fixed set of mutation types, most commonly defined by base substitution types (e.g., C>T) and trinucleotide contexts (e.g., TCT) for single-base substitutions (SBSs). By exploiting characteristic mutation patterns and variation in their frequencies across samples, this framework has proven powerful for disentangling mutational processes and inferring the mutational history of cancer genomes across tissues [10–12]. By construction, however, mutational signature analysis does not consider where in the genome those mutations occur, discarding potentially rich information about the interplay between mutational processes and their genomic context.

In the traditional formulation, therefore, mutational signatures reflect only the local effects of DNA sequence context. Yet mutation rates vary widely across the genome under the influence of additional genetic and epigenetic factors. At the *meso*-scale (spanning tens of base pairs), DNA secondary structure and protein–DNA interactions influence mutability [13–20]. Transcriptional activity and transcription-coupled nucleotide excision repair (TC-NER) modulate mutational patterns and regional mutational rates at the kilobase scale, while replication timing and chromatin organization do so on the order of megabases [13, 21–23]. Together, these factors induce substantial heterogeneity across the genome: megabase-sized regions can vary by more than five-fold in their mutation rates. Beyond affecting overall mutation rates, these processes also generate DNA strand-dependent mutations, as the transient exposure of single-stranded DNA during transcription and replication creates asymmetric opportunities for damage and enzymatic activity [13, 21, 23–25].

Understanding how these factors influence mutational processes could elucidate the mechanisms by which mutations arise. However, disentangling process-specific effects remains a challenge because mutational processes are rarely observed in isolation. A further challenge is extreme data sparsity in inferring a process’s genomic profile: even highly mutated cancer genomes contain only tens of thousands of mutations distributed across billions of possible sites.

Existing approaches address only limited aspects of these challenges. Non-negative matrix factorization (NMF)-based signature methods discover processes *de novo* by separating their spectra but do not model locus-level variation [26, 27]. Conversely, models of regional mutation rates and mutational hotspots capture variation across loci, but they neither deconvolve mutational processes nor explain inter-sample differences [28–33]. TensorSignatures jointly infers processes and their associations with genomic annotations, but requires coarse-grained discretization that lifts mutations from their native loci. This method also assumes mutation rates vary multiplicatively and independently with respect to epigenetic factors [34]. Finally, *ad hoc* strategies that first assign mutations to signatures and then correlate their genomic distributions with features [13, 14, 23, 35] are confounded by nonlinear and collinear associations between processes and covariates, and require *a priori* knowledge of the mutational spectra a process produces. As a result, despite extensive efforts, the distribution of mutational processes across the genome remains incompletely characterized.

Mutational signatures provide a static summary of how mutational processes affect the genome, but the outcomes of mutagenesis are dynamic – the intensity of process activity varies with genomic state and cellular conditions. Capturing this dynamic behavior requires a richer descriptive framework: mutational topography, defined as the joint variation in mutation spectra and genome-wide mutation rate profile. To this end, we developed MuTopia (Mutational Topography Inference and Analysis), a generative probabilistic framework that models mutagenesis as a joint function of mutation type, genomic locus, and sample-specific process activity, inferring mutation spectra and their genomic distributions simultaneously rather than sequentially. Applied across 15 tumor types, MuTopia reveals that mutational processes organize into conserved topographic archetypes shaped by damage, repair, replication, and chromatin state. Processes traditionally considered singular often comprise state-dependent forms that are distinguishable only through their genomic distributions.

## Results

### A generative model that learns the principles shaping mutational topography

MuTopia is a probabilistic generative model that, like NMF [36], decomposes mutational data into components representing distinct processes. Unlike these methods, MuTopia models not only each process’s mutational spectrum but also its genomic topography, parameterizing locus-dependent mutation rates as expressive, nonlinear functions of genomic and epigenetic covariates (**Figure 1a,b**). Concretely, the model generates each mutation by first selecting a process, then selecting a locus and mutation type according to that process’s topographic distribution. This defines a joint model that enables simultaneous inference of the active signatures and their genome-wide landscapes from sequencing data.

**Figure 1:**
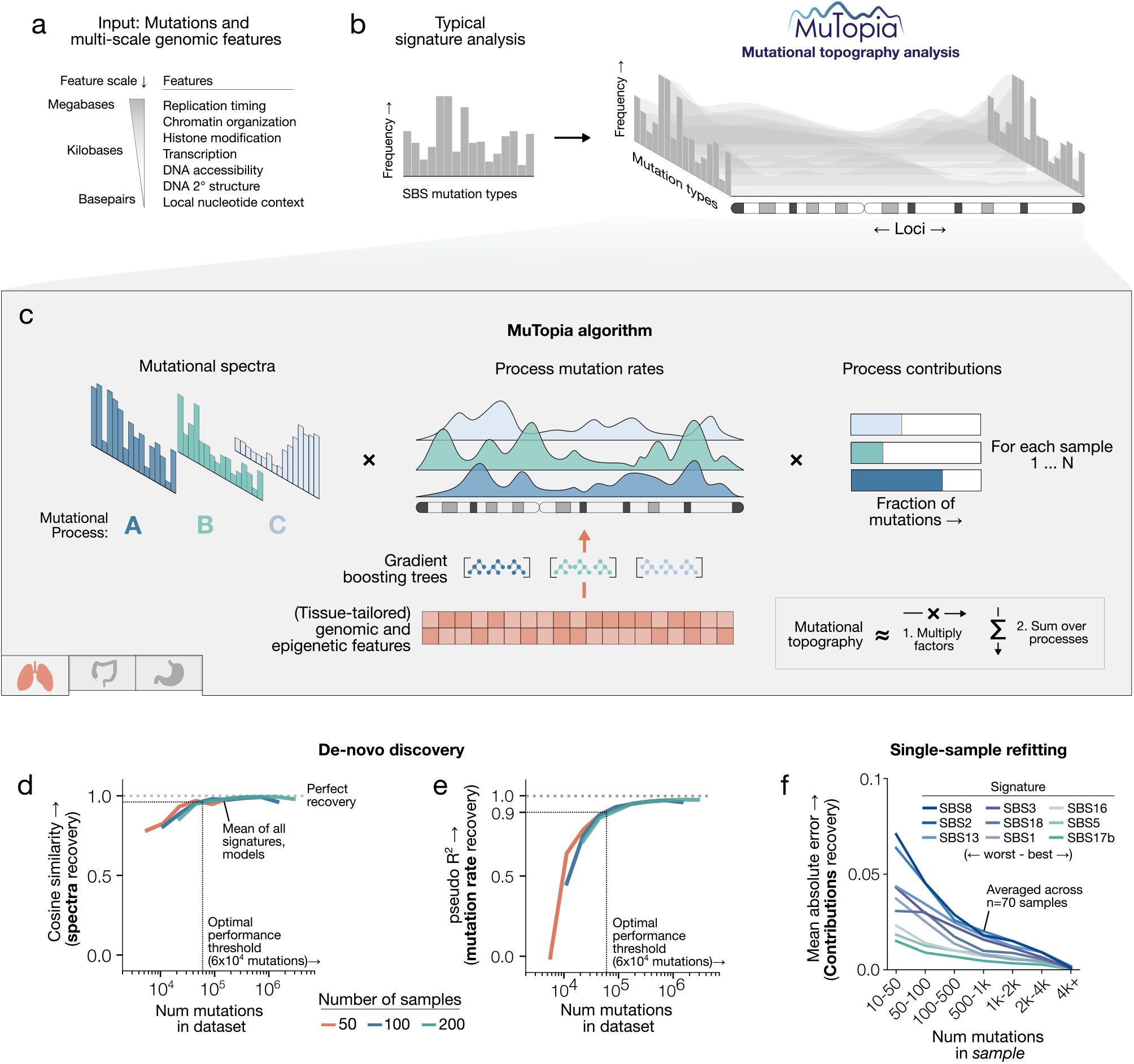
MuTopia model overview and benchmarking. **a)** Genomic features and their length scales used by MuTopia. **b)** Input data for standard signature analysis versus MuTopia topography analysis. **c)** Schematic of MuTopia’s factorization. Process mutation rates are learned as nonlinear functions of tissue-matched genomic features. **d)** Spectral signature recovery (cosine similarity) in simulated data across mutation burdens and cohort sizes (50, 100, 200 samples). **e)** Genome-wide mutation rate recovery (pseudo-*R*^2^) for the same simulations. **f)** Mean absolute error of single-sample exposure refitting across 70 simulated samples.

MuTopia addresses the high dimensionality of mutational topographies and the sparsity of mutation data through a structured factorization scheme. Each mutational process is decomposed into three latent components: a mutation type spectrum, a genome-wide mutation rate distribution (**Figure 1c**), and a strand-dependent adjustment term, regularized to favor uniform strand effects over mutation-type-specific asymmetries (**Supplementary Figure 1**). We model process-specific topographies as the product of these components, scaled by sample-level activity (“exposures”) (**Figure 1c**).

To capture pervasive nonlinear relationships between genomic features and mutation rates, MuTopia fits gradient-boosted tree regressors that predict each locus’s expected mutation count conditioned on genomic state (**Figure 1c, bottom**) [37]. By defining mutation rates based on features instead of estimating them independently for each genomic bin, we reduce the dimensionality of the model while yielding interpretable mappings between mutational processes and their genomic determinants. In addition, rather than using coarse, fixed-size bins, MuTopia utilizes a dynamic genome-binning strategy to integrate diverse genomic covariates spanning scales from base pairs to megabases. MuTopia first partitions the genome into 10-kilobase bins, which are then subdivided by fine-grained annotations such as gene bodies, ATAC-seq peaks, or DNA secondary structures (like hairpin loops) for mutation counting (**Supplementary Figure 2**). Since the model learns rates as functions of features, it is highly robust to sparsity, enabling the modeling of mutation rates at high resolution and across genomic scales spanning orders of magnitude.

Together, these design elements enable MuTopia to accurately recover high-dimensional mutational topographies from sparse data. We evaluated this capability through benchmarking on realistic tumor topographies simulated from a trained breast cancer model across a range of mutation burdens and cohort sizes (**Methods; Supplementary Figure 3a**). MuTopia accurately recovered both mutational spectra (cosine similarity; **Figure 1d**) and genome-wide mutation rate distributions (Poisson pseudo-*R*^2^ [38]; **Figure 1e**), with process recovery performance saturating at approximately 60,000 mutations per cohort (**Supplementary Figure 3b,c**). Since average per-sample mutation counts in prostate, breast, lung, and melanoma cohorts are approximately 3, 5, 30, and 100 thousand, respectively (**Supplementary Figure 4a,b**), this places most cancer cohorts within MuTopia’s accurate inference regime. Even at low mutational burdens, MuTopia could infer genome-wide rate distributions from as few as 2–10 samples given known signature spectra (**Supplementary Figure 3d**). For application to individual tumors, we implemented a refitting framework that estimates sample-level exposures from predefined topographic signatures with known spectra and genome-wide distributions (**Figure 1f**; **Supplementary Figure 3c**), allowing our collection of topographic signatures to be applied to new datasets without retraining.

### MuTopia accurately reconstructs genome-wide mutational topography across scales

We applied MuTopia to 15 tumor types from the PCAWG dataset and an additional compendium of breast cancer datasets [39], spanning wide ranges of sample sizes (42–760 tumors) and total mutation burdens (100,000 to 10 million mutations per dataset). For each tumor type, we constructed standardized, tissue-matched genomic feature sets from ENCODE (**Supplementary Methods; Supplementary Table S1**), compiled a dataset of mutations, optimized hyperparameters, and trained a MuTopia model (**Methods; Supplementary Figure 5**). We tested whether learned relationships between mutational processes and genomic context generalize beyond observed loci by evaluating model predictions using 3-fold cross-validation on held-out chromosomes. Chromosome 2 was withheld from both hyperparameter optimization and training across all tumor types, and used as a final test set (**Supplementary Figure 6a**). We reserved chromosome 2 for testing because of its large size and diversity of epigenetic states.

We contrasted MuTopia with alternatives that represent the range of currently available methods (**Supplementary Figure 6b**), including locus-only mutation rate models, sample-level signature models (for example, NMF-like approaches), and an adaptation of MuTopia that uses linear models rather than gradient-boosted tree regressors. Across all tumor types, MuTopia achieved superior reconstruction fidelity relative to these ablated models, as assessed by the pseudo-*R*^2^ metric (**Supplementary Figure 6c**).

In held-out genomic regions, MuTopia’s predictions of overall mutation rate showed strong concordance with observed data across tumor types, as illustrated for lung, breast, and stomach adenocarcinomas (**Figure 2a–c**, top subpanels). Beyond accurately predicting aggregate mutation rates, MuTopia recapitulated intricate strand-resolved shifts in mutational spectra across the genome (**Figure 2a–c**, middle subpanels). Here, we plotted the predicted mutation rate across loci for each of the 192 stranded single-base substitution contexts defined by trinucleotide sequence, clustered within each substitution type (C>T, G>A, etc.). This view of mutational topography revealed striking variation in mutational outcomes across tissues driven by differences in the active mutational processes, but also across loci within the same tumors. The ability to infer coherent, locus-specific spectral variation in genomic regions not used during training indicates that the learned signature–feature relationships capture generalizable dependencies between genomic context and mutational outcomes. Despite tumor type-specific differences in epigenomic state and mutational process activity, shared structure emerged across cancers: recurrent high- and low-mutability regions demarcated by pronounced spectral shifts. Transitions between mutagenic regimes corresponded with changes in gene density, chromatin state, and replication timing (**Figure 2a–c**, bottom subpanels), showing that genome state drives the coordinated modulation of both mutation rate and the local mutational spectrum. At finer genomic resolutions, the influence of gene expression and transcription-strand asymmetry on mutational topography became increasingly apparent (**Supplementary Figure 7**).

**Figure 2:**
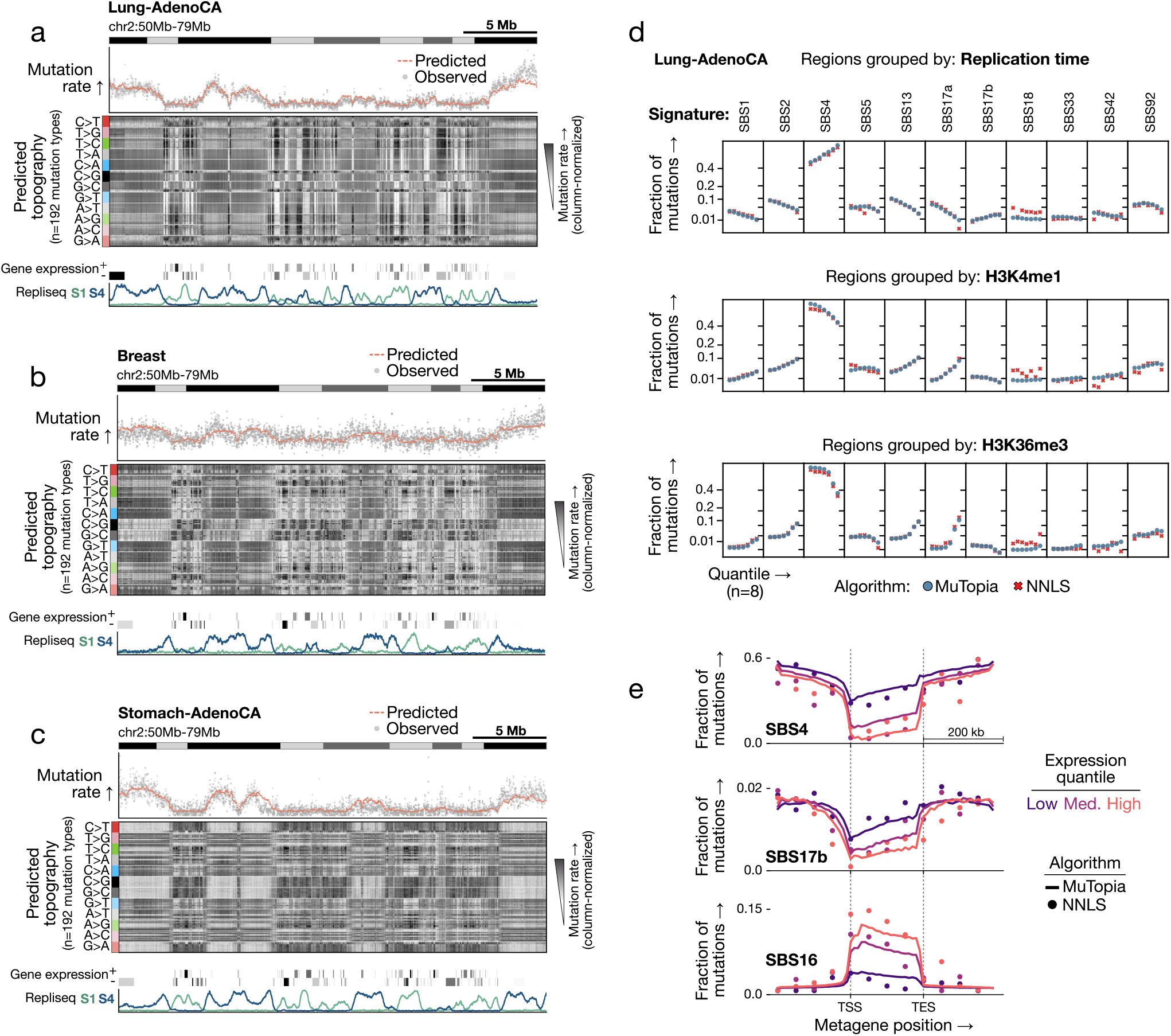
MuTopia captures multidimensional trends in mutational topography. **a)** Lung adenocarcinoma. Top: observed versus predicted marginal mutation rates. Middle: predicted mutational topography across 192 mutation types (column-normalized). Bottom: representative genomic features. **b,c)** Same as (a) for breast and stomach adenocarcinomas, respectively. **d)** Signature-specific mutation fractions across equal-size quantiles of the genome comparing MuTopia predictions to NNLS estimates, stratified by selected genomic features. **e)** Same as in (d), except along gene bodies. Genes were resized to align the transcription start site (TSS) and transcription termination site (TES) before binning.

As validation, we compared MuTopia’s signature-specific topographies to estimates obtained by non-negative least squares (NNLS) refitting of mutations aggregated across loci with shared genomic properties. MuTopia-predicted signature distributions showed strong concordance with NNLS-derived estimates across genomic features (**Figure 2d**) and along gene bodies (**Figure 2e**). For instance, exogenous damage processes were depleted within genes due to the activity of transcription-coupled NER, while SBS16 peaked near transcription start sites in proportion to gene expression. TC-NER can be error-prone when nearby lesions interfere with gap filling of the excised DNA patch, promoting the recruitment of translesion synthesis (TLS) polymerases [40]. The enrichment of SBS16 in highly transcribed regions is therefore consistent with TLS-mediated mutagenesis. The predicted shifts in mutational topography across gene-dense, highly transcribed regions (**Figure 2a–c**, middle subpanels) reflect coordinated reductions in damage-driven processes through TC-NER and corresponding increases in transcription-linked mutagenesis, illustrating how MuTopia disentangles the effects of overlapping mutational mechanisms.

Leveraging MuTopia’s ability to model genomic features at high resolution, we next asked whether this framework could reveal previously unrecognized structure in mutational topography. We identified a mutational process in microsatellite-stable breast tumors that matched the COSMIC signature SBS34 — which currently has no known etiology — and that was enriched within gene bodies and microsatellite regions (**Supplementary Figure 8**). SBS34 shares a similar mutational spectrum with a clustered signature described by Supek *et al*. [41]. Together with its enrichment near microsatellite regions, which can form secondary structures that impact transcription and replication, this suggests that SBS34 may reflect the activity of an error-prone polymerase operating over microsatellite tracts, potentially during the repair of transcribed sites.

These observations show that MuTopia accurately captures mutation topography consistent with empirical patterns, while its feature-based representation of genomic loci enables the generation of mechanistic hypotheses about the origins of mutational signatures.

### MuTopia elucidates the factors governing genomic profiles of mutagenesis

By deconvolving the contributions of individual mutational processes and linking them to genomic features, MuTopia enables direct interrogation of how mutagenic mechanisms interact with genome state. We illustrate this using stomach adenocarcinoma as a representative case study. We decomposed the overall mutation rate into signatures and inferred the genomic mutation rate profiles for each (**Figure 3a–c**). Then, we computed Shapley [42] value feature explanations for each signature to infer how changes in genome state influence its activity (**Figure 3d**; **Supplementary Figure 9**). Oxidative damage–associated signatures (SBS18, SBS17a, SBS17b) formed a coherent topographic group enriched in late-replicating heterochromatin and depleted from transcriptionally active regions (**Figure 3e**) [43, 44]. The uncharacterized signature SBS8 also grouped with this cluster. Its elevation in NER-deficient tumors implicates global genome-NER substrates as its source [45], and its C>A and T>A-rich spectrum is reminiscent of known oxidative damage signatures [3], together suggesting an endogenous oxidative etiology. Within this group, replication timing emerged as a key distinguishing feature, with subtypes of both SBS17b and SBS8 showing a shift toward earlier-replicating domains.

**Figure 3:**
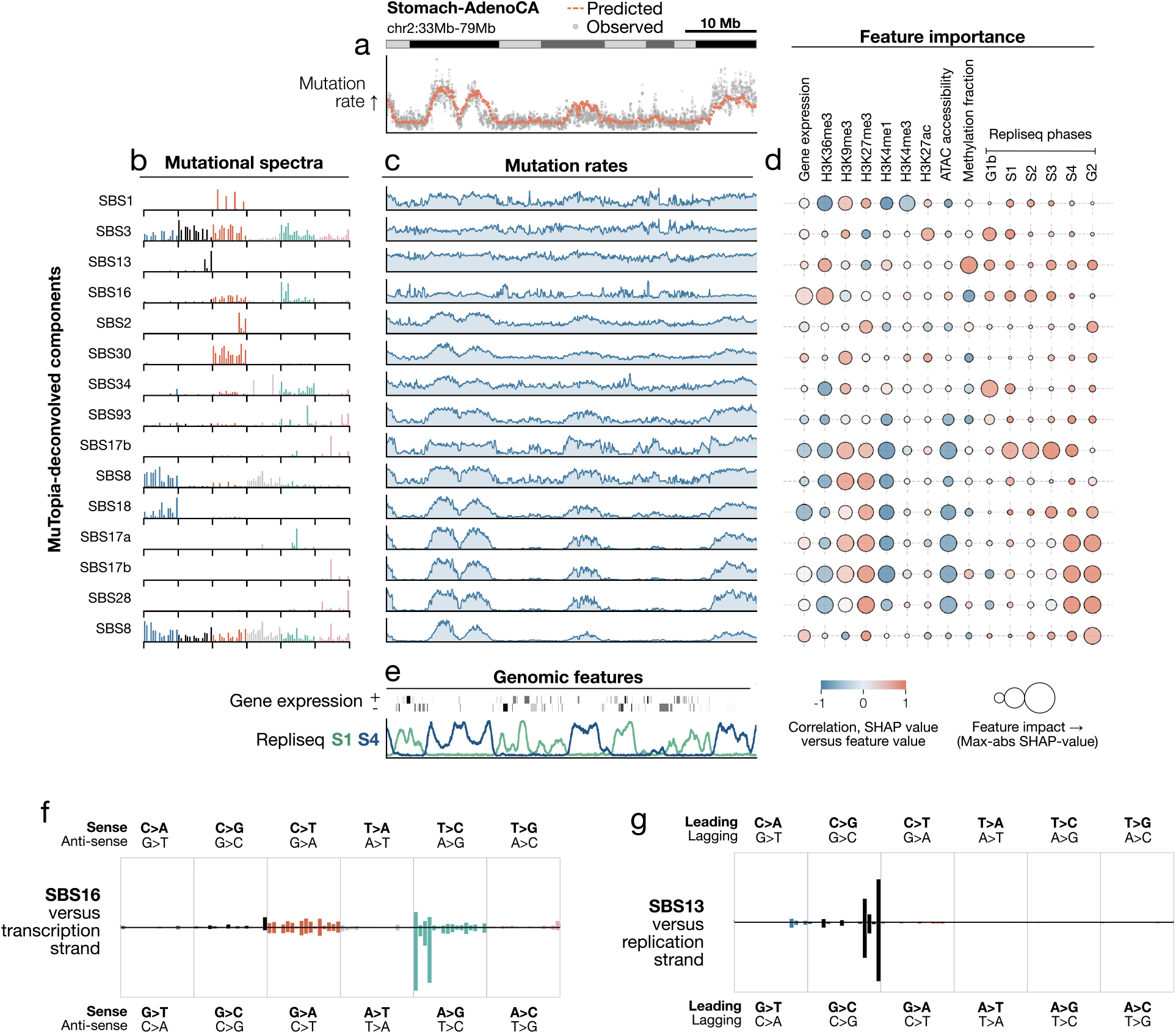
MuTopia decomposition of mutational processes in stomach adenocarcinomas. **a)** Observed versus predicted marginal mutation rates on held-out chromosome 2. **b)** Mutational spectra of discovered signatures, ordered by topographic clustering and named by COSMIC match. **c)** Predicted genome-wide mutation rates by signature. **d)** Shapley-derived feature associations per signature. Marker color: correlation between feature values and Shapley values. Marker size: 97^th^ percentile absolute Shapley value (feature impact). **e)** Representative genomic features driving variation at this resolution. **f)** Predicted mutational spectrum versus transcription strand for SBS16. **g)** Predicted mutational spectrum versus replication strand for SBS13.

Critically, topographic analysis resolved repair-dependent modes of mutagenesis that were indistinguishable from spectra alone. SBS8 separated into two spectrally similar but topographically distinct subtypes: one enriched in earlier-replicating regions (S1–S4) and in homologous recombination–deficient (HRD) tumors (classified using CHORD [46], p=9.5 × 10^−4^, Mann–Whitney U-test), and a second confined to late-replicating regions (S4–G2) in repair-proficient tumors (**Supplementary Figure 10**). In contrast, the canonical HRD signature SBS3 showed a pronounced bias toward very early-replicating regions (G1b–S1), consistent with replication-associated mutagenesis. Strand asymmetries provided additional resolution: SBS16 showed strong transcription-strand bias proportional to gene expression (**Figure 3f, Supplementary Figure 11a**), while SBS13 showed pronounced lagging replication strand bias with early replication enrichment (**Figure 3g**; **Supplementary Figure 11b**), in contrast to the late-replicating bias of SBS2.

Together, these patterns illustrate how genomic topography refines the interpretation of mutational processes beyond spectral similarity, revealing mechanistic stratification within spectrally similar signatures and recurrent mutation rate profiles shaped by replication dynamics and DNA repair.

### A pan-cancer analysis reveals topographic archetypes of somatic mutagenesis

To identify unifying principles of mutational topography, we sought to compare mutation rate profiles learned across tumor types. Because each model leveraged tissue-specific epigenomic features, however, the inferred genome-wide mutation rate profiles naturally reflected the chromatin and replication landscapes of their respective cell types. We resolved this issue by re-estimating the rate profiles using a shared epigenomic reference — here, lung tissue, chosen for the breadth and quality of its ENCODE feature tracks — thereby isolating intrinsic topographic properties of mutational signatures independent of tissue-specific genomic state (**Figure 4a**; **Supplementary Table S2**). We then focused our analysis on 208 out of 220 signatures that either exhibited consistent spectra across multiple tumor types or matched established COSMIC reference signatures (**Supplementary Figures 12, 13**; **Supplementary Table S3**). The remaining signatures either reflected known artifact-associated patterns or exhibited mixed spectra consistent with multiple underlying processes, limiting their interpretability.

**Figure 4:**
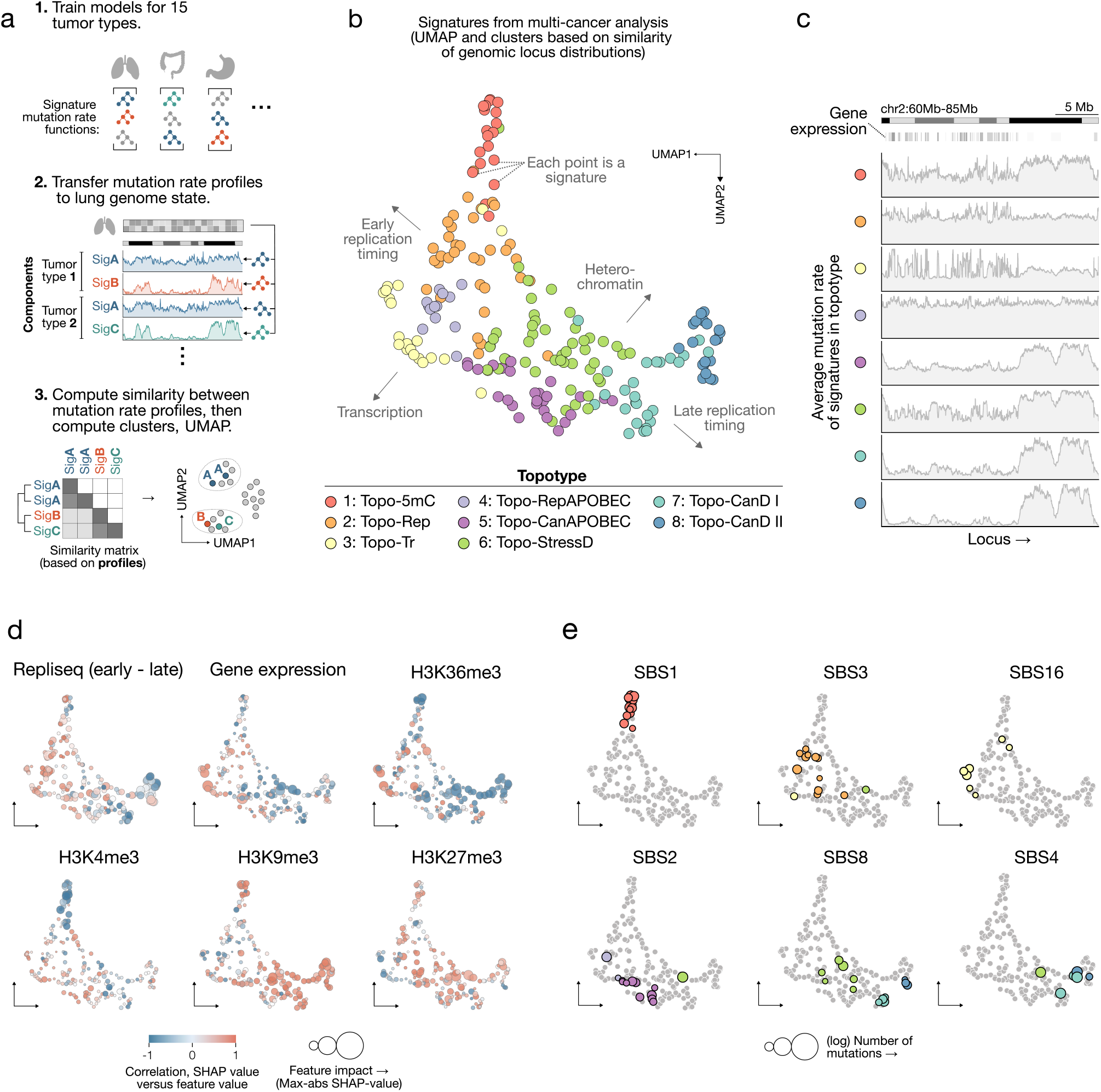
Pan-cancer analysis of mutational processes’ genomic profiles. **a)** Steps of the unified analysis. **b)** UMAP of 208 signatures from 15 tumor types, clustered by similarity of genomic mutation rate profiles using the Leiden algorithm and manually annotated into topotypes. **c)** Average mutation rate profiles for each topotype. **d)** Shapley-derived feature associations. Marker color: correlation. Marker size: effect size. Repli-seq values are computed as early (G1b + S1 + S2) minus late (S4 + G2b). **e)** Selected COSMIC SBS annotations highlighted on the UMAP. Marker size indicates log-transformed mutation count.

By clustering their genome-wide mutation rate profiles, we identified eight groups of processes with distinct locus-dependent patterns (**Figure 4b,c**; **Supplementary Figure 14a**; **Supplementary Table S4**). These clusters represent conserved archetypes of mutagenesis shared across processes and tumor types, which we refer to as topography types, or “topotypes”. UMAP projection revealed that the clusters were structured primarily by replication timing and chromatin state: early-replicating, euchromatin-associated topotypes were separated from late-replicating, heterochromatin-associated topotypes in the embedding (**Figure 4d**; **Supplementary Tables S5, S6**). Spectrally related signatures from independent tumor types frequently clustered within the same topotypes (**Figure 4e**; **Supplementary Figure 14b,c**), consistent with the idea that common mutational processes produce similar locus-dependent mutation patterns. We leveraged this co-clustering, together with known mechanisms of well-characterized mutational processes, to infer the biological basis of these topotypes.

One topotype (cluster 1) was defined by signatures acting preferentially at methylated cytosines. SBS1, which represents C>T mutations at 5-methylcytosine (5mC) CpG sites arising from spontaneous deamination or polymerase *ϵ*–associated errors [10, 47], mapped to this topotype across all tissues (**Figure 4e**). Signatures in this cluster were enriched in regions marked by H3K27me3 and H3K9me3 and strongly depleted from active gene regulatory domains marked instead by H3K36me3 and H3K4me3 (**Figure 4d**), consistent with the localization of 5mC to heterochromatin and its depletion at hypomethylated CpG islands in active promoters. This cluster also contained a C>G mutational process at CpG sites that resembled COSMIC SBS98, a more recent addition to the COSMIC catalog. This mutational pattern has been reported to result from active CpG demethylation involving oxidized 5mC and abasic-site–mediated base-excision repair, observed in both germline and cancer genomes [26, 33, 48]. Therefore, we refer to this cluster as the 5mC-associated topotype (Topo-5mC). The grouping of distinct but mechanistically related processes within a single cluster demonstrates that MuTopia links mutational processes operating on a shared genomic substrate.

In contrast to the repressive, methylated chromatin associated with Topo-5mC, cluster 3, which we term the transcription-associated topotype (Topo-Tr), comprised signatures enriched in transcriptionally active regions and within highly expressed gene bodies exhibiting a strong transcription-strand bias. In addition to SBS16, this cluster contained SBS34, SBS41, and the colibactin signature SBS88, several of which share SBS16’s enrichment in AT-rich trinucleotide contexts (**Supplementary Figure 15**). While the mutational signatures of Y-family polymerases are not fully defined, the observed enrichment in AT-rich contexts across many signatures — drawing parallels to polymerase *η*–associated mutagenesis — may reflect their activity in the context of translesion synthesis during transcription-coupled repair.

Damage-associated mutational processes — including endogenous damage by SBS8, tobacco-related SBS4, oxidation-linked SBS18 and SBS17a/b, and UV-associated SBS7 and SBS38 — fell within three adjacent topotypes (clusters 6, 7, and 8) (**Figure 4e**; **Supplementary Figure 14b,c**). Clusters 7 and 8, dominated by oxidative damage processes, occupied the late-replicating, heterochromatic region of the embedding space (**Supplementary Figures 16, 17**) and defined the canonical topographic regime of damage-associated mutagenesis. These clusters were distinguished by the degree of constitutive heterochromatin confinement. SBS18 and SBS36 preferentially aligned with the broader late-replication profile of cluster 7 (Topo-CanD-I, or “Canonical Damage”), while SBS17 signatures were enriched in the more constitutive heterochromatin–confined cluster 8 (Topo-CanD-II). In contrast, cluster 6 exhibited a systematic shift toward earlier replication timing (S1–S3) and a stronger depletion of mutations in regions marked by active chromatin features and high gene expression. In addition to UV-associated signatures, it contained variants of oxidative damage signatures (SBS8, SBS4, SBS18, and SBS17) that co-occurred with their canonical counterparts within the same tumor types but showed distinct topographic profiles. The localization of UV-associated damage signatures (SBS7a/c/d, SBS38) within cluster 6 suggests that lesions arising from distinct sources — such as bulky UV-induced photoproducts versus smaller oxidative base modifications — are processed by repair pathways with different spatial and temporal dynamics that give rise to distinct genomic topographies.

Notably, we observed two distinct SBS8 subtypes co-occurring within the same cohort across multiple tumor types, recapitulating our earlier observations in stomach adenocarcinomas and showing that SBS8 consistently splits between topotypes Topo-CanD-I/II and cluster 6. Further analysis confirmed that this split reflects HR status: in HR-proficient (HRP) tumors, SBS8 adopts the canonical damage-like profile of Topo-CanD-I/II, whereas in HRD tumors, it adopts the cluster 6 mutation rate profile (**Figure 5a, Supplementary Figure 18a,b**). We therefore designate cluster 6 as the stress-associated damage topotype (Topo-StressD), reflecting its links to replication stress and repair deficiency–modulated damage processing. Signatures adopting distinct genomic distributions depending on HR status demonstrate that mutational topography reveals intrinsic cellular stress states.

**Figure 5:**
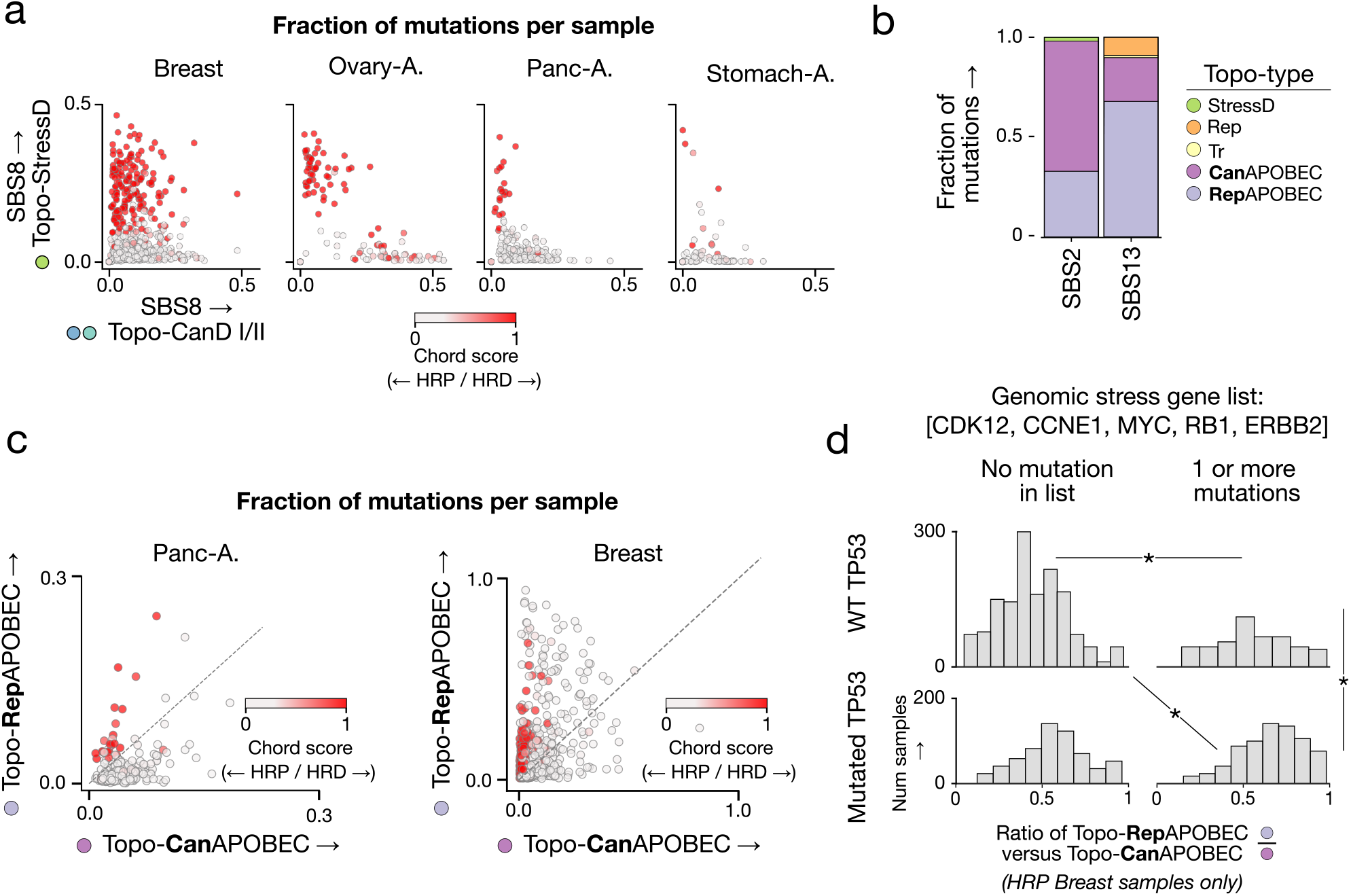
HRD and replication stress induce topotype shifts. **a)** SBS8 mutations attributed to Topo-StressD versus Topo-CanD-I/II topotypes across four tumor types, colored by CHORD HRD likelihood. **b)** Fraction of SBS2 and SBS13 mutations in each topotype across all datasets. **c)** APOBEC mutations attributed to TopoRepAPOBEC versus Topo-CanAPOBEC topotypes in pancreatic and breast tumors, colored by CHORD HRD likelihood. **d)** Ratio of Topo-RepAPOBEC to Topo-CanAPOBEC mutagenesis in HRP breast tumors, stratified by replication stress gene alterations and TP53 status. (Mann–Whitney U-test, *: adj. p *<* 0.05)

Cluster 2, which we refer to as the replication-associated topotype (Topo-Rep), showed weak associations with chromatin features and gene expression, but a preference for early replication timing (G1b–S2 phase). This cluster contained the canonical HRD signature SBS3, as well as a C>G-enriched component (Mu1), both enriched in HRD relative to HRP tumors. In tumor types where Mu1 was detected, it separated from the broader SBS3-like distribution due to its distinctive replication timing profile (**Supplementary Figure 18c,d**). While both components exhibited early-replicating enrichment characteristic of Topo-Rep, Mu1 showed a more pronounced shift toward very early-replicating regions, particularly in the G1b phase. Together with SBS8, these processes were enriched in HRD tumors, indicating that HRD-associated mutagenesis comprises multiple co-occurring processes with separable topographic profiles. Leveraging this decomposition, MuTopia’s topography-aware analysis enables more robust classification of ovarian HRD tumors than mutational spectra alone (**Supplementary Figure 18e**).

Finally, the remaining two clusters (4 and 5) were both enriched for APOBEC-associated signatures but showed clearly distinct topographies. While cluster 5 was biased toward late-replicating regions and predominantly contained the signature SBS2, cluster 4 exhibited a flat mutation rate profile akin to Topo-Rep and was enriched for SBS13 (**Figure 5b**). We thus termed cluster 5 “Topo-CanAPOBEC” (canonical APOBEC) and cluster 4 “Topo-RepAPOBEC” (replication-associated APOBEC). Since replication stress-induced fork-stalling events can expose single-stranded DNA to APOBEC editing, we used HRD as a proxy to test how this might influence the topography of APOBEC mutagenesis. Strikingly, in pancreatic adenocarcinoma tumors, HRD-mediated replication stress induced a shift from primarily Topo-CanAPOBEC to Topo-RepAPOBEC (**Figure 5c**).

In breast tumors, we observed three APOBEC subtypes: SBS2 in Topo-CanAPOBEC, and a variant of both SBS2 and SBS13 in Topo-RepAPOBEC (**Supplementary Figure 19a**). HR status was again a key factor in determining the rate of Topo-CanAPOBEC versus Topo-RepAPOBEC activity, but it was insufficient to explain all Topo-RepAPOBEC-high tumors. Replication stress can also arise from rapid entry into S phase triggered by oncogenes [49–51], which may cause replication-transcription collisions that lead to replication fork stalling. We compiled a set of genes known to induce replication stress through distinct mechanisms when amplified (CCNE1, MYC, and ERBB2) or lost (CDK12 and RB1). We found that replication stress, defined by the alterations in these genes, led to an enrichment in Topo-RepAPOBEC activity in HR-proficient breast tumors when paired with mutations in TP53 (**Figure 5d**; **Supplementary Figure 19b**). The higher abundance of SBS13 versus SBS2 in Topo-RepAPOBEC is consistent with the known roles of UNG and REV1 in generating C>G mutations, linking SBS13 to REV1 activity during replication [52]. These results suggest that replication stress changes the genomic profile of APOBEC mutagenesis, increasing deamination activity and UNG- and REV1-mediated error-prone repair in early-replicating regions.

Together, these results indicate that mutational topography converges upon a small number of conserved archetypes primarily governed by replication timing, chromatin state, and DNA repair activity.

### The landscape of mutational topotypes across cancers

To summarize the joint dependencies between mutational processes and their genomic distributions, we quantified topotype attributions for each signature in each tumor type (**Figure 6a**; **Supplementary Table S7**). While some signatures — SBS1, SBS3, SBS16 — maintained largely the same topotypes across tissues, others, particularly damage-associated processes, exhibited substantial variability. This representation makes explicit how spectrally similar signatures can manifest diverse genome-wide profiles depending on cellular state.

**Figure 6:**
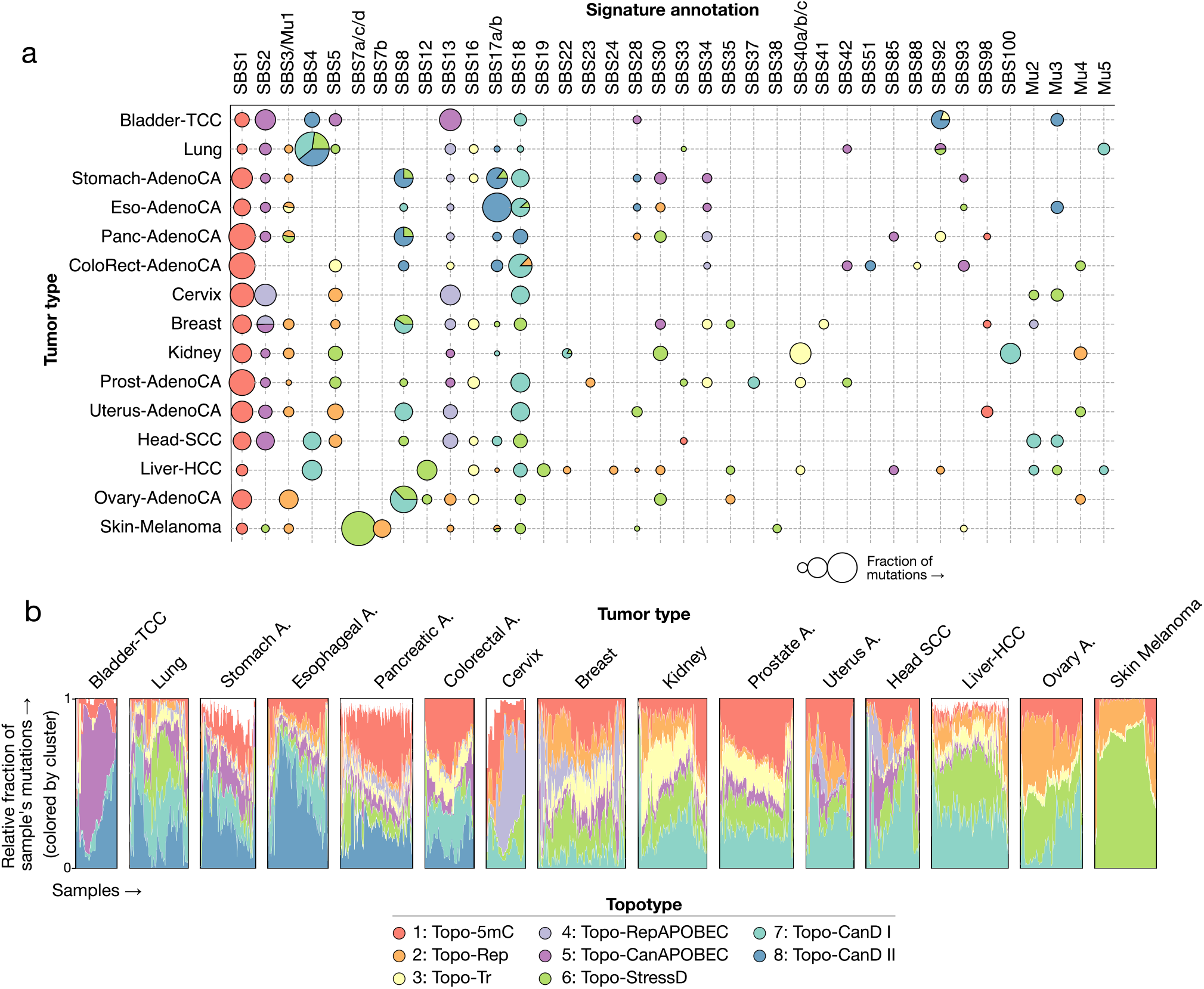
The pan-cancer landscape of mutational topography. **a)** Mutations attributed to a given topotype are shown for all signatures across tumor types. The marker size indicates the relative fraction of mutations attributed to that signature within a given tumor type. **b)** The topotype exposures per sample across tumor types. Exposures for unannotated signatures are shown in white.

At the sample level, topotype mixtures showed clear patterns of inter-tumor heterogeneity (**Figure 6b**). In breast and ovarian cancers, topotype composition recapitulated HRD-defined molecular classes. Even in tumor types dominated by a single mutational process, such as lung cancers with smoking-associated damage, we observed pronounced diversity in topotype composition, highlighting the potential of topographic modeling in resolving patient-specific variation in DNA repair capacity and replication competency.

Alongside this inter-tumor diversity, we observed surprising concordance across tumor types. Despite being subject to diverse sources and intensities of DNA damage — like oxidation in stomach and esophageal tissues versus UV damage in skin — the relative balance of damage, replication, and transcriptional topotypes was remarkably consistent across tumor types (**Supplementary Figure 20**). This suggests that shared constraints on the proficiency and fidelity of repair pathways primarily determine the genomic distribution of mutations, not the biases and properties of the instigating DNA damage.

This landscape view of mutational topography provides a genome-resolved, cell-type-agnostic framework for understanding somatic mutagenesis. By linking mutational processes to their genomic distributions, it extends mutational signature analysis beyond nucleotide spectra to enable comparative analysis of mutational mechanisms, tumor states, and environmental exposures across cancers.

## Discussion

Mutational signatures have historically been defined by their mutational spectra alone. Our results demonstrate that this representation is incomplete. Mutational processes possess characteristic genomic topography, or topotypes, that are conserved across cancers, shaped by replication timing and chromatin state, and altered by cellular repair proficiency. While the genomic distribution of some processes directly reflects the substrate on which damage occurs — as in the heterochromatic confinement of 5mC-associated mutagenesis — the convergence of diverse damage sources onto shared topotypes suggests that, for many processes, repair pathway engagement and replication dynamics, rather than damage propensity alone, are the principal determinants of where mutations ultimately accumulate [53, 54].

This topographic representation revealed biological features unseen through traditional mutational signature analysis, highlighting that mutational processes are not static but can manifest with distinct, state-dependent topographies. Several damage-associated processes (SBS4, SBS8, SBS17, and SBS18) exhibit subtypes that co-occur within the same tumor types and share mutational spectra, yet differ in their genome-wide distributions, including shifts across replication domains and differential associations with gene expression. These differences are consistent with modulation by repair defects and replication stress, as illustrated by SBS8, which separates into canonical and replication stress–associated topographies enriched in HRD tumors. Apart from damage processes, replication stress systematically reshapes the topography of APOBEC editing, influencing both its spectrum and mutation rate profile. Together, these findings refine conventional signature definitions and demonstrate that mutational processes are best understood as context-dependent programs rather than fixed spectral entities.

By integrating the genomic state directly into a generative framework, MuTopia unifies the analysis of mutational spectra and mutation rates. We redefine mutational signatures to incorporate genome-wide topographic structure, moving beyond purely spectral representations. To connect our work to the wider field, we aligned MuTopia-derived signatures with COSMIC nomenclature based on spectral similarity. Notably, joint *de novo* discovery enabled the decoupling of previously composite signatures in multiple instances. Processes traditionally considered singular, such as HRD-associated mutagenesis (SBS3), comprise subtypes with distinct genomic distributions. This extends to the clock-like SBS5, which separates into components with replication- and damage-associated C>T enrichment and transcription-associated T>C enrichment, most closely matching COSMIC SBS30 and SBS16, respectively (**Supplementary Figure 21**). These observations suggest that widely observed signatures may comprise multiple underlying processes that are separable only through their genomic topographies.

The COSMIC database provides topographic associations for some signatures [35], but these were calculated one feature at a time. Since multiple genomic features — replication timing, chromatin state, and gene expression — jointly shape topography, single-feature analyses cannot accurately define the isolated effect of each feature to determine which are the primary drivers of a signature’s genomic distribution. Prior *post-hoc* approaches that first assign mutations to signatures and then examine their genomic distributions share this limitation [13, 55], and cannot detect cases where the same spectral signature adopts different topographic programs in different cellular contexts. TensorSignatures jointly inferred mutational spectra and genomic state coefficients [34], but it retained a linear model structure and pre-defined categorical feature bins. In contrast, MuTopia models mutation rates as expressive nonlinear functions of tissue-matched genomic features, with Shapley-value attribution isolating each feature’s marginal contribution. Strand biases are similarly decomposed as an explicit model component, enabling transcription- and replication-strand effects to be disentangled from each other and from the confounding influence of other genomic features.

Beyond the conceptual advances revealed by our pan-cancer analysis, MuTopia surpasses existing approaches in several key methodological aspects. First, it adopts a generative, probabilistic framework that enables principled modeling of mutation counts and uncertainty. Second, it integrates genomic features across hierarchical genomic scales, capturing diverse biological processes ranging from local sequence context to large-scale chromatin and replication domains. Third, it models the effects of genomic state through nonlinear functions, using gradient-boosted trees with Shapley value–based attribution to provide both flexibility and interpretability. By linking mutational processes to interpretable genomic features, topography-aware decomposition enables more precise attribution of previously uncharacterized signatures to their underlying etiologies. Finally, MuTopia effectively addresses the sparsity of mutational data, enabling robust inference even in low-mutation or small-cohort settings. The resulting pan-cancer collection of topographic signatures, available through open-source software, provides a resource for investigating mutagenesis across tissues. MuTopia’s refitting method enables its application to new datasets and tumor types.

Several limitations remain. Our formulation does not explicitly distinguish DNA damage induction from repair outcomes, and we did not attempt to decouple these into separate model components. Repair pathways operate at distinct cell-cycle stages and interact nonlinearly, rendering a fully identifiable decomposition infeasible without controlled experimental systems involving single-pathway perturbations — an important future direction. We also relied on high-quality ENCODE normal tissue epigenomes for modeling; however, epigenetic and replication landscapes change during tumor progression, so our feature space may not fully represent mutational profiles later in tumor evolution. The topotypes we identified may be sensitive to the features used for mutation rate modeling. However, since MuTopia’s predictions appear quite concordant with observed mutation rates (**Figure 2a–c**), we expect that the addition of more features will only marginally reduce the residual variation and, therefore, will not substantially influence the results. Nevertheless, MuTopia is readily adaptable to incorporate additional genomic features of interest. Finally, although we focused here on single-base substitutions, the framework is general and can be extended to other classes of somatic variation, including indels and structural variants.

In sum, our results establish mutational topography as a fundamental dimension of somatic genome evolution and provide a framework for understanding how genome organization shapes the mutational landscape of human cancers.

## Methods

### MuTopia: a generative model of mutational topography

We model somatic mutagenesis as a mixture of *K* latent mutational processes. For each sample, let 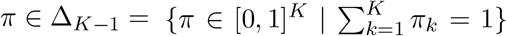 denote the vector of (normalized) process exposures, where *π*_*k*_ is the fraction of mutations attributed to process *k*. We place a Dirichlet prior on these exposures, *π* ∼ Dir(*α*), with 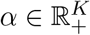.

To generate a mutation in a sample, one first draws a mutational process *z* | *π* ∼ Cat(*π*), so that *p*(*z* = *k* | *π*) = *π*_*k*_, and then draws a genomic bin *b* = 1, …, *N*_bins_ and mutation type *m* = 1, …, *N*_types_ from the process-specific distribution *p*(*b, m* | *z* = *k, θ*). Here, bins are feature-defined groups of genomic loci (see below), mutation types correspond to the 192 trinucleotide-by-base-change categories, and *θ* denotes the model parameters determining the process-specific topographies. Repeating this procedure yields the somatic mutations observed in each sample.

Let 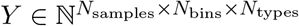 denote the mutation counts, where *Y*_*nbm*_ is the number of mutations of type *m* observed in sample *n* in genomic bin *b*. Under this generative model, the marginal likelihood is

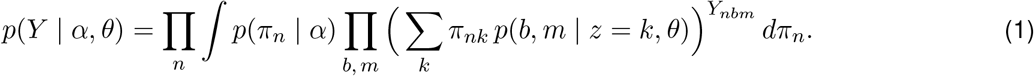

(**Supplementary Methods**). The distribution *p*(*b, m* | *z* = *k, θ*) defines the mutational topography of process *k*. Because mutation data are far too sparse to estimate this distribution independently for each genomic bin, we instead model it as a function of genomic features. Specifically, let 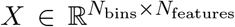 denote the matrix of genomic features, with *X*_*b*_ denoting the feature vector for genomic bin *b*. We decompose the process-specific topography into two log-additive functions of genomic state: the macro-scale effects 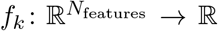, which capture variation in the overall mutation rate across the genome, and the spectra effects 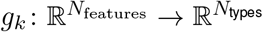, which capture local variation in the mutational spectrum (**Figure 1c**). Together with the context-availability factor *t*_*bm*_, which accounts for the number of sites in bin *b* at which mutations of type *m* can occur, we define:

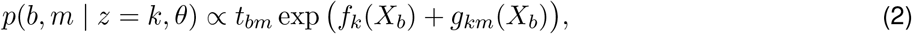

where ∝ denotes normalization over all bins and mutation types, and *θ* denotes the collection of parameters defining the functions *f*_*k*_ and *g*_*k*_ (**Supplementary Methods**).

#### Macro-scale effects

We model the macro-scale effects *f*_*k*_ using histogram gradient-boosted tree (HGBT) regression [37] on genomic features expected to influence the overall mutation rate across the genome. These features include replication timing, histone modifications, chromatin accessibility, and gene expression (**Supplementary Methods, Table 3**). Using HGBT regression allows MuTopia to capture nonlinear relationships between genomic state and mutation rate that are pervasive in mutation data (**Supplementary Methods**).

#### Spectra effects

The function *g*_*k*_ extends the usual notion of a mutational signature. Instead of assigning each process a single fixed mutation spectrum, MuTopia allows the spectrum of process *k* to vary with local genomic context. Our motivation is that different classes of genomic features may affect mutagenesis in different ways. Macro-scale features are expected to primarily influence a process’s mutation rate profile, for example, through differences in overall repair activity. In contrast, meso-scale features—such as gene bodies, microsatellite regions, or CTCF-binding sites—may interact more directly with DNA damage and repair factors, and can therefore locally alter the relative frequencies of mutation types a process produces. Accordingly, for each process *k*, the model combines a baseline mutation spectrum with sparse *ℓ*_1_-penalized adjustments driven by meso-scale and strand-oriented features through a linear function *g*_*k*_, which we fit using a Poisson regression objective (**Supplementary Methods**). This encodes the assumption that we expect most meso-scale features do not change a process’s mutational spectrum, and when they do, they are more likely to produce small changes rather than the wholesale reshaping of the signature.

#### Genomic bins and features

MuTopia accommodates genomic features spanning scales from base pairs to megabases through a flexible binning scheme. To define genomic bins, we first partition the genome at every position where a discrete feature – either macro- or meso-scale – changes state, and introduce additional breakpoints at a configurable fixed-width interval (10 kb by default). This yields a set of fine-grained segments within which the discrete genomic state is constant. We then aggregate segments with identical discrete feature states into bins, even when those segments are not contiguous in the genome. For continuous features, we assign to each bin the average feature value across all bases in that bin. MuTopia, therefore, groups loci by shared genomic state rather than by physical continuity, enabling joint modeling of small features and broader macro-scale regions.

For each tumor type, we constructed tissue-matched reference sets of genetic and epigenetic features using an automated procedure that selected the highest-quality unperturbed ENCODE experiment for each assay and tissue of origin (**Supplementary Methods; Supplementary Table S1**). We selected features that were both available across all target tissues and likely to influence mutation rates. The final set of features included: Gene expression and strandedness; DNase-seq chromatin accessibility signal; ATAC-seq peak locations; fraction of methylated CpGs via bisulfite sequencing; DNA GC content; ChIP-seq signal for the histone modifications H3K27ac, H3K4me1, H3K4me3, H3K36me3, H3K27me3, and H3K9me3; Repli-seq signal for the phases G1b, S1, S2, S3, S4, and G2; and replication strand (**Supplementary Methods, Table 3**).

### Model outputs

For downstream analysis, we consider the following model quantities. In the definitions below, samples are weighted in proportion to their burden, i.e., *p*(*n*) ∝∑_*b,m*_ *Y*_*nbm*_ when computing cohort-level quantities.

1. **Mutational spectra:** The spectrum of process *k*, given by *p*(*m* | *z* = *k, θ*) = ∑_*b*_ *p*(*b, m* | *z* = *k, θ*).
2. **Process genomic distributions:** The genomic distribution of process *k* across bins, given by *p*(*b* | *z* = *k, θ*) = ∑_*m*_ *p*(*b, m* | *z* = *k, θ*).
3. **Process contributions:** The normalized exposure of process *k* in sample *n*, given by *π*_*nk*_.
4. **Per-sample topographies:** The mutational topography of sample *n*, given by *p*(*b, m* | *θ, n*) = ∑_*k*_ *π*_*nk*_ *p*(*b, m* | *z* = *k, θ*).
5. **Genomic mutational spectra:** The cohort-level mutational spectrum at genomic bin *b*, given by *p*(*m* | *b, θ*) ∝∑_*n,k*_ *p*(*n*) *π*_*nk*_ *p*(*b, m* | *z* = *k, θ*).
6. **Normalized mutation rate:** The cohort-level normalized mutation rate, given by *p*(*b* | *θ*) = ∑_*n,k*_ *p*(*n*) *π*_*nk*_ *p*(*b* | *z* = *k, θ*).
7. **Feature explanations:** We compute Shapley values [56] for each process’s macro-scale effects *f*_*k*_, yielding 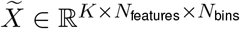. For each process *k* and feature *i*, we report the “feature impact” as Quantile 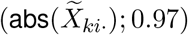 and the “directionality” as 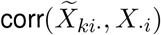. Intuitively, feature impact captures the extent of variation in mutation rates induced by a feature, while the directionality encodes the strength of the linear relationship between feature values and rates.

### Model training

#### Variational inference

We use mean-field variational inference in the style of Latent Dirichlet Allocation [57, 58] to fit the model. Specifically, because the integrals over the exposures *π* are analytically intractable, we estimate the model parameters (*α, θ*) and approximate posterior distributions over the latent variables (*π, z*) by maximizing the evidence lower bound (ELBO), a tractable lower bound on the log marginal likelihood in (1). To this end, we introduce the factorized variational distribution *q*(*π, z* | *β, ϕ*) = Π_*n*_ *q*(*π*_*n*_ | *β*_*n*_) Π_*b,m*_ *q*(*z*_*nbm*_ | *ϕ*_*nbm*_), where the variational parameters *β*_*n*_ parameterize approximate Dirichlet posteriors over the exposures, and *ϕ*_*nbm*_ parameterize approximate posteriors over mutation-to-process assignments. Optimization proceeds by coordinate ascent: in the variational E-step, we update *β* and *ϕ* given the current model parameters, and in the M-step, we update *α* and *θ* given the current variational parameters. Intuitively, the E-step updates the approximate posterior over the sample-specific exposures and mutation-to-process assignments under the current model, and the M-step updates the process topographies and prior parameters using those variational estimates.

#### Stochastic optimization and scalability

To scale model training to datasets containing millions of mutations, we use stochastic variational inference with subsampling over genomic bins rather than samples. This choice has a practical advantage: when updating the topographies from subsampled bins, we aggregate information across all samples, which yields less noisy regression targets and improves training stability. Similarly, because the variational parameters for all samples are updated at each iteration, the Dirichlet prior *α* can be updated using information from the entire cohort at every step. We implemented MuTopia using just-in-time compilation via *numba* [59] to further speed up model training. Full derivations and optimization details are provided in the Supplementary Methods.

### Estimating model outputs

The model outputs defined above depend on the topography parameters and latent exposures. During training, MuTopia learns topography parameters 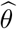, together with variational approximations to the posterior distributions of the sample-specific exposures *q*(*π*_*n*_ | *β*_*n*_). We therefore estimate the model outputs using 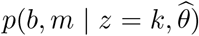 and 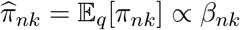.

### Model evaluation

#### Pseudo-*R*^2^

To assess the goodness of fit on held-out data, we use a pseudo-*R*^2^ metric based on the multinomial log-likelihood [38]. Let *Y* denote the held-out mutation counts. For any model that yields per-sample topography estimates 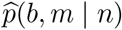 over genomic bins and mutation types, we define the score

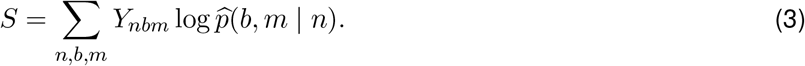

We compare this score to those of a null model and a saturated model. For the null model, we use the context-availability distribution 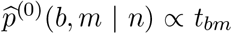, with corresponding score *S*^(0)^. For the saturated model, we use the empirical distribution of the held-out data 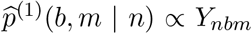, with corresponding score *S*^(1)^ . We then define the pseudo *R*^2^ as

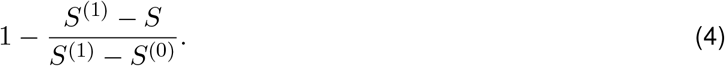

By construction, this normalization assigns a pseudo-*R*^2^ of 0 to the null model and 1 to the saturated model. Since no estimated model can outperform the saturated model, pseudo-*R*^2^ ≤ 1.

#### Model ablation benchmark

We used pseudo-*R*^2^ to compare the full MuTopia model with the following ablated models:

- **MuTopia-Linear:** Identical to MuTopia, except the nonlinear macro-scale effects *f*_*k*_ were replaced with a linear model.
- **LDA:** The macro-scale effects were replaced with constant terms, *f*_*k*_ = 0, and the spectra effects were restricted to the baseline factors alone. This makes the model equivalent to LDA, up to the inclusion of the context availability factor.
- **Locus only:** The number of mutational processes was fixed to *K* = 1, removing the ability to model sample-specific variation and reducing the distribution across genomic bins to a gradient-boosted tree regression. The spectra effects were restricted to the baseline factors alone, such that the resulting signature spectrum equals the empirical mutational spectrum across the entire dataset.

For each tumor type, we computed pseudo-*R*^2^ after holding out one of chromosomes 1–3 for evaluation.

#### Simulation benchmark

To evaluate MuTopia’s ability to recover latent mutational processes from limited data, we first trained the model on the large breast cancer cohort from [39], yielding reference parameters *α, θ*, and *π*_*n*_. We selected seven components resembling known COSMIC signatures and used their inferred topographies as the basis for simulation. For each synthetic sample, we drew an exposure vector *π*_*n*_ by sampling uniformly from the set of inferred exposures in the original cohort and sampled ⌊*γM*_*n*_⌋ mutations distributed across processes according to *π*_*n*_, where *M*_*n*_ is the number of mutations in the corresponding original sample and *γ* ∈ ℝ_+_ controls the overall mutation burden. This procedure preserves realistic covariance between processes and burdens.

For each combination of *N*_samples_ ∈ {50, 100, 200} and *γ* ∈ {2^−4^, 2^−3^, …, 2^4^}, we trained three replicate models with *K* = 7 and selected the best-fitting replicate by pseudo-*R*^2^. Recovery performance was assessed using cosine similarity between matched spectra and Poisson pseudo-*R*^2^ between matched rate profiles, the latter serving as the analog of correlation for comparing Poisson-distributed data.

#### Exposure refitting benchmark

To evaluate exposure estimation using a pretrained model, we selected breast cancer samples with more than 5,000 mutations and treated their MuTopia-estimated exposures 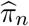 as ground truth. For each sample, we subsampled mutations at varying mutational burdens and re-estimated *π* while keeping the global model parameters *α* and *θ* fixed and optimizing only the sample-specific variational parameters. We report the mean absolute error between the exposures estimated from the subsampled and full data.

### Validation

#### NNLS validation

To validate MuTopia’s inferred topographies, we partitioned the genomic bins into groups based on individual genomic features. For epigenetic marks and replication timing, we used eight quantile-based groups, each containing the same number of base pairs. Coarser grouping was necessary to ensure sufficiently high mutation counts for stable non-negative least squares (NNLS) estimates. For gene expression, we divided bins into 15 groups based on their distance and orientation relative to the nearest gene, and then further subdivided these groups based on that gene’s expression level (low, medium, or high).

For each group *G* ⊆ {1, …, *N*_bins_}, we estimated the relative contribution of mutational process *k* using MuTopia based on

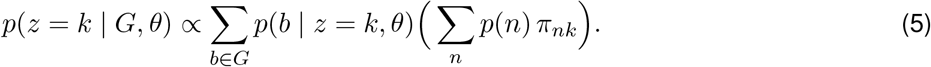

As a comparison, we pooled all mutations across samples within each group and applied NNLS to estimate mutational process proportions from the aggregate mutational spectrum.

#### HRD classification

To compare the predictive value of topography-aware versus spectral features for HRD classification, we trained *ℓ*_2_-penalized logistic regression models on 113 ovarian adenocarcinoma samples, with HRD labels predicted using CHORD [60], using two feature sets: (1) the MuTopia-inferred process contributions 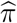 and (2) the 96-channel mutational spectrum. We selected the *ℓ*_2_ penalty via 5-fold cross-validation across samples using the complete whole-genome mutation data. To assess robustness to mutation burden, we then subsampled mutations from each sample, recomputed the features from the reduced data, and applied the trained logistic regression model to predict HR status.

### Pan-cancer analysis of topography archetypes

To start, we compiled datasets for 15 tumor types using mutation calls from the PCAWG consortium, pairing each tumor type with genomic features collected using the automated strategy outlined in the supplementary methods. Using the annotations of Jin *et al*. [26], we excluded 89 mismatch repair–deficient and hypermutated samples, as these exhibit mutation distributions that differ markedly from the remainder of the cohorts. We then trained MuTopia models for each tumor type separately, selecting the number of components *K* and other hyperparameters via random search with HyperBand [61] pruning, scored by pseudo-*R*^2^ on the held-out chromosome 2. To initialize each tumor type model, we seeded MuTopia with COSMIC signature spectra reported in that tumor type at a sample frequency greater than 2%, again following Jin *et al*..

We performed hyperparameter tuning in three rounds. Between rounds, we removed samples as outliers if the sample alone comprised the majority of mutations attributed to a rare component. In total, 11 of approximately 2,500 samples were removed during this iterative curation. In addition, after fitting, we removed initialized signatures that appeared only very rarely across samples under the learned model, retaining only signatures supported by sufficient activity.

After hyperparameter optimization and model selection, the final set of tumor-type models yielded 220 mutational components in total. To enable pan-cancer comparison of locus-dependent topographies, we transferred the learned mutation-rate profiles for each component—parameterized by the trained *f*_*k*_ ensembles—to a shared genome state by applying each component’s trained gradient boosting tree ensemble to the lung feature matrix, chosen for the breadth and quality of available ENCODE tracks. Lung-derived signatures were well-mixed within the resulting topotype clusters, indicating no strong reference tissue bias. We then computed pairwise cosine similarity between all transferred mutation-rate profiles across chromosome 2, which was held out from training in all tumor-type models during hyperparameter optimization (multi-fold validation showed the model exhibits minimal variance in performance dependent on the held-out chromosome). Chromosome 2 was selected for testing and analysis because it is large, epigenetically diverse, and exhibits copy number alterations at a lower rate than other chromosomes across most tumor types. From the cosine similarity matrix, we constructed a nearest neighbors graph, which served as the basis for both community detection via the Leiden algorithm [62] and two-dimensional embedding via UMAP [63]. For the initial Leiden clustering, we utilized a resolution of 0.2; for the UMAP projection, we used 10 nearest neighbors, a *min_dist* of 0.1, and a *negative_sampling_rate* of 3.

Finally, we performed manual curation to refine the Leiden clusters into coherent topotypes and to assign the best-matched COSMIC signature designations to each component. For the pan-cancer meta-analysis, we used 208 components (comprising 98.8% of the total mutations in the dataset) that met at least one of three criteria: (i) the component closely matched a known COSMIC signature; (ii) the component did not match any COSMIC signature but was independently discovered in three or more tumor types; or (iii) the component was observed in only one tumor type but was common across samples within that type (two components met this criterion). Of 220 total components, 12 were rejected under this filter, one of which was an instance of the known artifact signature SBS60.

### Sample ingestion

#### Mutation weighting

For each mutation in each sample, we recorded the genomic bin with which it intersected, its mutation type (one of 192 options), and a weight used to quantify its contribution to the local mutation rate. We then constructed *Y*_*nbm*_ as the sum of the weights of mutations in sample *n*, genomic bin *b*, and mutation type *m*. The total weight, then, should be proportional to the mutation rate at that locus. Because tumors experience frequent copy number alterations over the course of their evolution, however, we cannot simply use the mutation count as the weight directly – the number of mutations observed at a locus depends on both the total DNA available to mutate (proportional to copy number) and the cumulative time of mutagenic exposure at that copy number state.

Instead, we estimate the mutation rate for sample *n*, bin *b*, and mutation type *m, λ*_*nbm*_, using the fact that 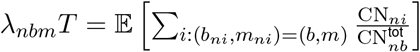, where (*b*_*ni*_, *m*_*ni*_) are the bin and mutation type of mutation *i* in sample *n*, CN_*ni*_ is the apparent copy number of the mutation, 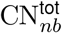 is the total copy number at that locus in the tumor at the time of sequencing, and *T* is a proportionality constant related to the time of exposure [64]. For a given mutation, the quantity 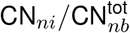 is related to its observed variant allele frequency (VAF_*ni*_) in a tumor sample by:

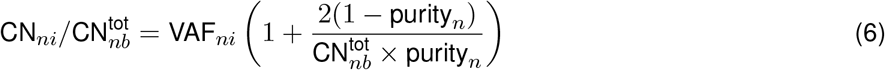

where VAF_*ni*_ is subject to sampling noise, and purity_*n*_ refers to the fraction of the sampled DNA belonging to tumor rather than non-malignant cells. Note that when 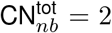, the equation above reduces to 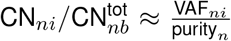 For simplicity, we take the weight of each mutation to be 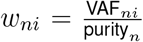 so that 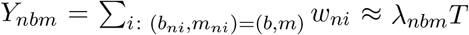 . This method enables the fast and robust estimation of normalized mutation rates across loci with variable copy number histories.

#### Clustered mutation adjustment

Clustered mutations — consecutive somatic mutations in close genomic proximity — frequently arise from processive or burst-like mutagenic activity at a single locus, such as APOBEC editing of exposed ssDNA or UV-induced tandem damage. Because such events reflect a single mutagenic opportunity rather than independent sampling of the genome-wide mutation rate, treating each clustered mutation as an independent observation would inflate the apparent local mutation rate and distort topography estimates. We define clustered mutations as those that are closer than expected under an inhomogeneous point process parameterized by the average background mutation rate across a cohort. If mutations are clustered, we divide their weights by the size of the cluster so that each group contributes the equivalent of one observation.

For each mutation *i* in sample *n*, we estimate the expected number of mutations at that base as 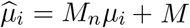, where *M*_*n*_ is the total number of mutations in the sample, *µ*_*i*_ is the relative number of mutations per base averaged over a 50-kb window around the mutation across the cohort, and *M* is a small pseudocount added to reduce the variance of the estimate (*M* = 1 by default). Under the null hypothesis *H*_0_ that mutations arose from a Poisson process with rate 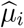, the distance *D*_*i*_ to the next mutation on the same chromosome (assuming a copy number of 2) is exponentially distributed, 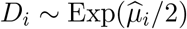. We reject *H*_0_ in favor of *H*_1_ — that mutation *i* lies closer to the next than the local Poisson rate predicts, and thus arises from a clustered event — at level *α* when 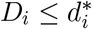, where the critical distance 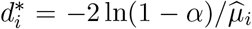 is the *α*-quantile of this distribution. Consecutive mutations on the same chromosome are then grouped into a cluster whenever their separation is at most 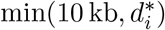. By default, we set *α* = 0.005. During inference of sample-level exposures, we retain the option to forgo cluster-size adjustment.

## Data availability

PCAWG data were downloaded from “https://www.synapse.org/#!Synapse:syn11726601/files/”. Signatures for model initialization were downloaded from Jin *et al*. [26]. All genome features were collected from the ENCODE data portal [65]. HRD likelihood scores were obtained from Nguyen *et al*. [46]. Sample-level driver mutation calls were obtained from Lee *et al*. [39]. The datasets we constructed for the pan-cancer modeling and analysis can be found at https://zenodo.org/records/18803136.

## Code availability

MuTopia is available as an open-source, Python-based analysis package and command line tool at https://github.com/sigscape/MuTopia.

## Acknowledgments

This work was supported by grants to PJP (NIH R01CA269805 and R01HG012573) and DCG (DoD OC230088) and to PJP/DCG from the Quadrangle Fund for Advancing and Seeding Translational Research (QFASTR) at Harvard Medical School.

## Author contributions

AL conceived of the project, implemented the model, performed data analyses, designed and compiled figures, and co-wrote the manuscript. SSL contributed to code development, tested the codebase, and edited the manuscript. JPH edited the manuscript. BG assisted in mathematical derivations, optimization implementation, and plot design, and co-wrote the methodology. HJ provided conceptual guidance and analysis advice. DG conducted the pan-cancer analysis, oversaw model implementation and training, designed figures, co-wrote the manuscript, and supervised the study. PJP supervised the study, provided conceptual guidance, and edited the figures and manuscript.

## Supplementary methods

### Overview of the probabilistic model

MuTopia is a latent-variable generative model for somatic mutations across genomic bins and mutation types. For each sample *n*, the model introduces a vector of normalized process exposures *π*_*n*_ ∈ Δ_*K*−1_, drawn from a Dirichlet prior, *π*_*n*_ ∼ Dir(*α*). Each mutation is then generated in two steps: first, a latent mutational process *z*_*ni*_ ∈ {1, …, *K*} is sampled from a categorical distribution with parameters *π*_*n*_, and second, a genomic bin *b*_*ni*_ and mutation type *m*_*ni*_ are sampled from the process-specific distribution *p*(*b, m* | *z*_*ni*_ = *k, θ*). The parameters *θ* define these process-specific topographies through the macro-scale functions *f*_*k*_ and spectra functions *g*_*k*_, together with the context-availability term *t*_*bm*_. In this way, MuTopia combines sample-specific mixture proportions with process-specific, feature-dependent mutation distributions across the genome.

### Mutation-level likelihood

Let *M*_*n*_ denote the number of mutations observed in sample *n* (the mutational burden). We write

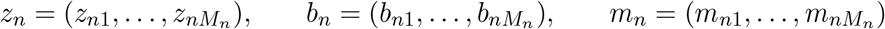

for the vectors of latent process assignments, genomic bins, and mutation types in sample *n*. Under the model, we have *p*(*z*_*ni*_ = *k* | *π*_*n*_) = *π*_*nk*_, and the complete-data likelihood for sample *n* therefore factorizes as

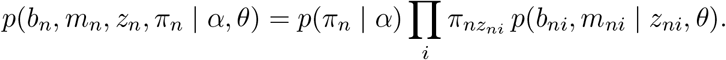

Marginalizing over the latent assignments *z*_*n*_ gives

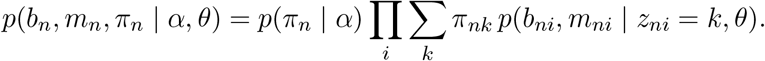

Finally, marginalizing over the latent exposures and taking the product over samples yields the marginal likelihood

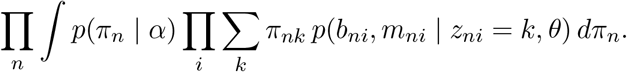

### Count-level likelihood

Now let

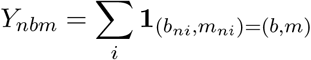

denote the number of mutations of type *m* observed in bin *b* for sample *n*. Since the mutation-level likelihood depends on the observations only through the counts of each bin–type pair in each sample, we can group identical terms to rewrite the marginal likelihood as

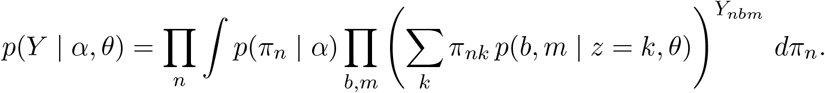

This is the same marginal likelihood as above, rewritten in terms of the aggregated counts *Y*_*nbm*_.

### Variational inference

Direct maximization of the marginal likelihood is intractable because it requires integrating over the latent exposures and summing over all mutation-to-process assignments. Variational inference addresses this by introducing a tractable family of distributions over the latent variables and deriving the evidence lower bound (ELBO), a lower bound on the marginal log-likelihood that can be computed efficiently. Maximizing the ELBO yields a practical surrogate objective for learning the model parameters (*α, θ*) while simultaneously approximating the posterior distribution over the latent variables [1].

We begin with a mean-field variational approximation at the mutation level,

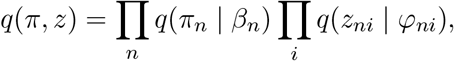

where *q*(*π*_*n*_ | *β*_*n*_) is Dirichlet and *q*(*z*_*ni*_ | *φ*_*ni*_) is categorical. For a single sample *n*, the ELBO is

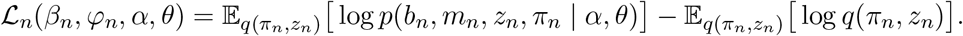

Summing over samples gives the full ELBO

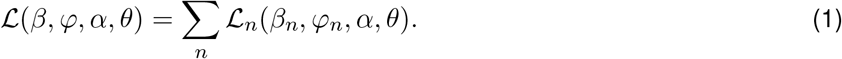

### Mutation-level ELBO

The explicit derivation of the ELBO is identical to that of latent Dirichlet allocation (LDA) [2], except that the vocabulary index is replaced here by the genomic bin–mutation-type pair (*b, m*). Using the mean-field variational family above, the per-sample ELBO can be written as

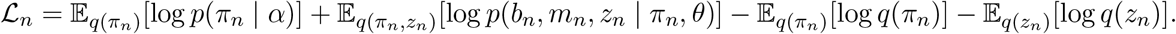

Since *q*(*π*_*n*_ | *β*_*n*_) is Dirichlet, the prior term is

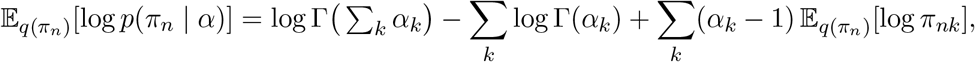

and the variational entropy term is

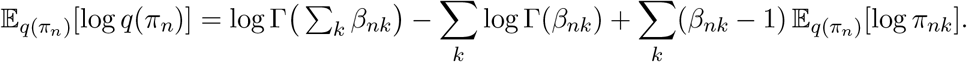

Here, Γ denotes the Gamma function, and the expectation can be computed explicitly using the digamma function *ψ* via

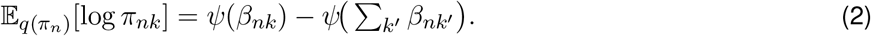

Next, we have

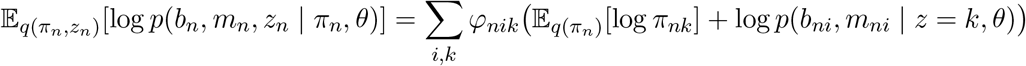

and

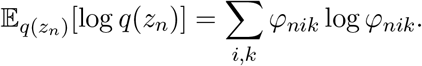

Combining these expressions and using

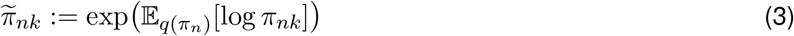

for notational convenience gives

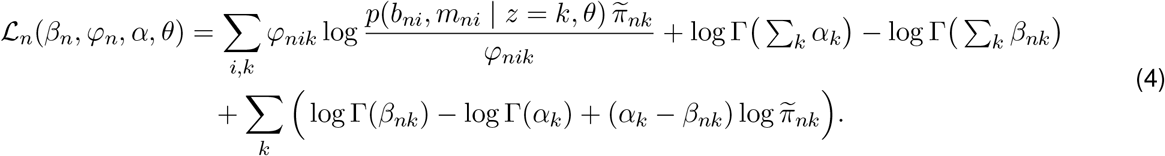

Summing over samples gives the full ELBO.

### Count-level ELBO

Because the likelihood depends on each mutation only through its observed bin–type pair (*b*_*ni*_, *m*_*ni*_), mutations in the same sample with the same pair (*b, m*) are exchangeable under both the model and the variational family. Consequently, the optimal variational parameters satisfy

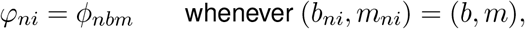

for some shared *ϕ*_*nbm*_ ∈ Δ_*K*−1_. Thus, although the variational family was introduced at the mutation level, the optimal variational distribution can be reparameterized using one probabilistic mutation-to-process assignment per sample, bin, and mutation type. Grouping the terms in (4) by (*b, m*) then yields

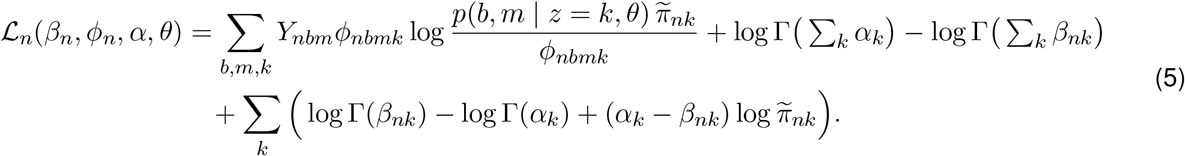

Summing over samples gives the full ELBO,

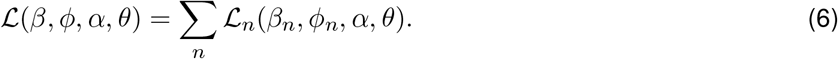

This is the same ELBO as above, rewritten in terms of the aggregated counts *Y*_*nbm*_. In practice, this representation is preferable because it avoids redundant variational parameters for mutations with identical observed bin–type pairs. We henceforth use the *ϕ*-parameterization of the model.

## Model training

MuTopia model training proceeds by maximizing (6) via coordinate ascent: in the variational E-step, we update *β* and *ϕ* given the current model parameters, and in the M-step, we update *α* and *θ* given the current variational parameters. Intuitively, the E-step updates the approximate posterior over the sample-specific exposures and mutation-to-process assignments under the current model, and the M-step updates the process topographies and prior parameters using those variational estimates.

### Optimizing the variational parameters

For fixed model parameters (*α, θ*), the variational E-step maximizes the ELBO with respect to the variational parameters *ϕ* and *β*. These updates are similar to LDA [2], except that the vocabulary index is replaced here by the genomic bin–mutation-type pair (*b, m*).

#### Updating *ϕ*_*nbmk*_

The terms in ℒ that depend on *ϕ*_*nbm*_ ∈ Δ_*K*−1_ are

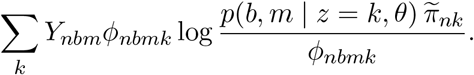

Using a Lagrange multiplier for the simplex constraint, the optimum satisfies

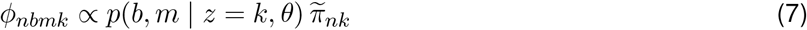

with normalization over *k*.

#### Updating *β*_*nk*_

The terms in ℒ that depend on *β*_*nk*_ are

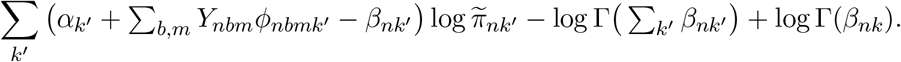

Optimizing over *β*_*n*_ yields the maximum at

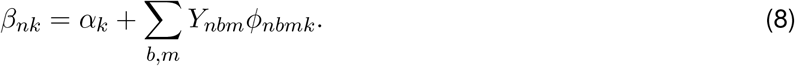

Thus, *β*_*nk*_ is given by the prior pseudo-count *α*_*k*_ plus the expected number of mutations in sample *n* assigned to process *k*.

The variational E-step therefore consists of alternating the updates in (7) and (8) until convergence.

### Optimizing the model parameters

#### Updating *α*

The terms in ℒ that depend on *α*_*k*_ are

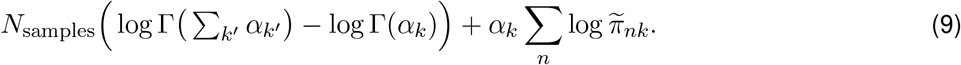

To optimize *α*, we follow [2, A.4.2] and use a Newton–Raphson algorithm with derivatives

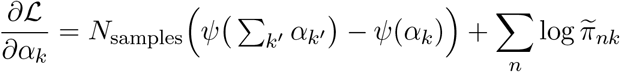

and Hessian

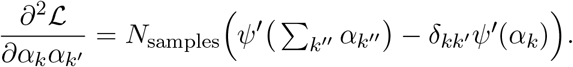

#### Updating *θ*

The terms in ℒ that depend on *θ* are

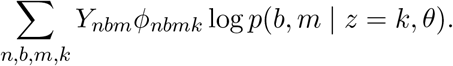

Under any natural parameterization of the model, the topography parameters are process-specific rather than shared across mutational processes. Accordingly, we write *θ* = (*θ*_1_, …, *θ*_*K*_), where *θ*_*k*_ denotes the parameters governing the topography of process *k*. Optimizing ℒ with respect to *θ* is therefore equivalent to optimizing ℒ with respect to each *θ*_*k*_ separately. For fixed *k*, we define the signature-assigned “soft counts”

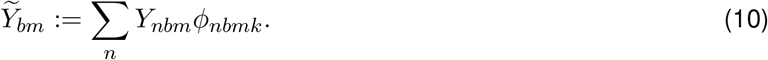

The terms that depend on *θ*_*k*_ can then be written as

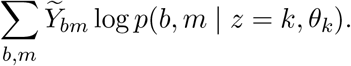

Thus, for each process *k*, the M-step reduces to maximizing a multinomial log-likelihood in which 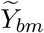 acts as the expected number of mutations assigned to process *k* in bin *b* and mutation type *m*.

For any unnormalized mutation rate *µ*_*bm*_ = *µ*_*bm*_(*θ*_*k*_) *>* 0, write the process-specific topography in the generic form

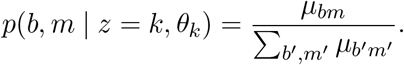

Substituting this into the objective gives

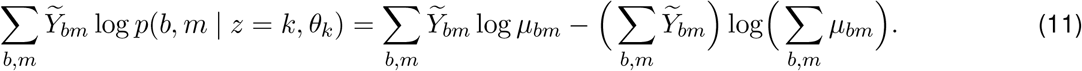

Direct optimization of this multinomial objective is inconvenient because the probabilities must remain normalized over all bin–type pairs, so the objective is not separable in *µ*_*bm*_. To avoid this, we apply the multinomial–Poisson transformation [3]. Introduce an auxiliary scalar parameter *c*_*k*_ ∈ ℝ, define

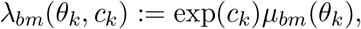

and consider the Poisson objective

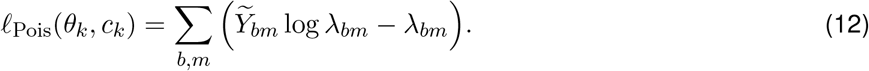

Substituting *λ*_*bm*_ = exp(*c*_*k*_)*µ*_*bm*_ yields

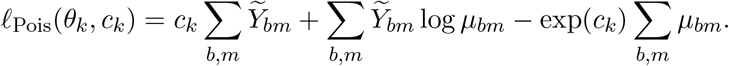

Maximizing over *c*_*k*_ gives

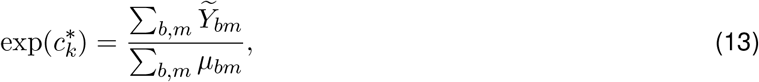

and substituting this back into the Poisson objective yields

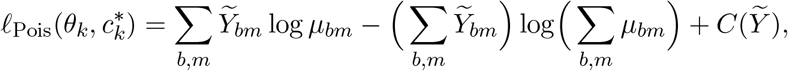

where 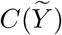 is a constant that depends only on 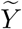. Comparing this expression with (11), we see that the profiled Poisson objective is equal to the multinomial objective up to an additive constant. Hence both objectives have the same maximizer in *θ*_*k*_. Therefore, updating *θ*_*k*_ is equivalent to maximizing *ℓ*_Pois_(*θ*_*k*_, *c*_*k*_) with respect to both *θ*_*k*_ and *c*_*k*_, and then retaining the optimal 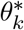.

In MuTopia, the unnormalized mutation rate is

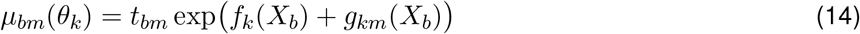

so that

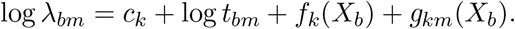

Thus, the M-step for *θ*_*k*_ can be reformulated as maximizing a Poisson log-likelihood with offset log *t*_*bm*_, soft count response 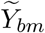, and process-specific intercept *c*_*k*_. The key advantage of this reformulation is that it replaces the normalized multinomial objective by a loss that is separable across bin–type pairs. In practice, this allows *f*_*k*_ and *g*_*k*_ to be fit using any model class that can be trained against a Poisson loss, including flexible estimators such as gradient boosted trees.

#### Updating the macro-scale effects *f*_*k*_

Let 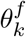 denote the parameters determining the macro-scale effects of process *k*. As shown in the previous section, maximizing ℒ with respect to 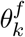 is equivalent to maximizing the Poisson objective

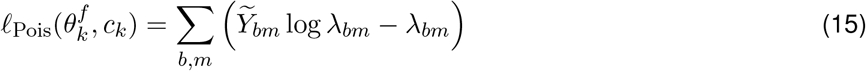

with respect to both 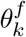 and *c*_*k*_, and then retaining the optimal 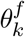. Here,

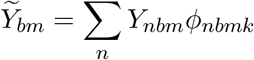

denotes the soft count assigned to process *k* in bin *b* and mutation type *m*, and

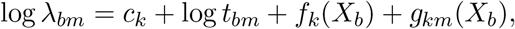

where *g*_*k*_ is treated as fixed throughout this update.

To optimize this objective with respect to 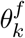 using standard machinery, it is convenient to rewrite it as a weighted Poisson loss whose rates depend on 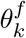 only through *f*_*k*_. Since the macro-scale effects do not vary with mutation type, it is also natural to aggregate over mutation types. For fixed *c*_*k*_, define

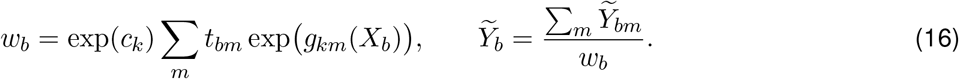

Then

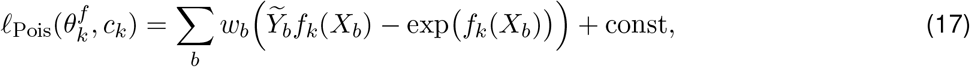

where const denotes terms independent of 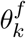. Thus, for fixed *c*_*k*_, updating *f*_*k*_ reduces to maximizing a weighted Poisson log-likelihood over bins, with sample weights *w*_*b*_, responses 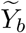, and predictors *X*_*b*_. This form of the loss allows us to fit *f*_*k*_ using the scikit-learn [4] implementation of histogram gradient-boosted tree (HGBT) regression [5] with a Poisson objective. In practice, when updating *f*_*k*_, we restrict *X*_*b*_ to the macro-scale features 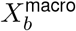 (**Table 3**).

For fixed 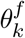, the optimal *c*_*k*_ is given by the general update from (13). With the process topographies parameterized as in MuTopia, this reads

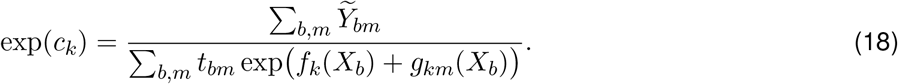

The updates for 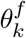 and *c*_*k*_ are alternated until convergence.

#### Incremental tree growth

Rather than fitting a new boosted tree model for *f*_*k*_ from scratch at every iteration, we instead maintain a single growing ensemble and append only a small number of additional trees at each update. This substantially improves training speed. We use this strategy both within the alternating updates of 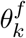 and *c*_*k*_, and across successive iterations of the outer training loop over all variational and model parameters.

Specifically, let 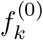 denote the current estimate of the macro-scale effects. To learn an additive update *f*_*k*_, we optimize the Poisson objective (15) under the parameterization

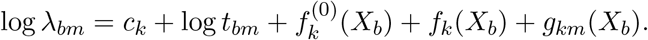

Thus, the newly learned function *f*_*k*_ is interpreted as an increment to the current estimate 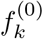. As before, for fixed *c*_*k*_ and 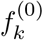, the objective can be rewritten as a weighted Poisson loss. The only difference is that the weights must now also absorb the contribution of the current macro-scale estimate. Define

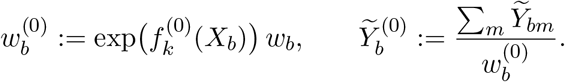

Then

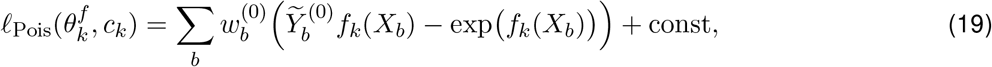

where const denotes terms independent of the additive update *f*_*k*_. We fit this update using HGBT regression with a small number of boosting stages, and then update the macro-scale effects via

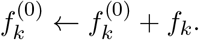

Repeated application of this procedure yields a single additive ensemble for the macro-scale effects, rather than a sequence of independently refit models.

#### Updating the spectra effects *g*_*k*_

Let 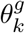 denote the parameters determining the spectra effects of process *k*. As shown above, maximizing ℒ with respect to 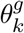 is equivalent to maximizing the Poisson objective

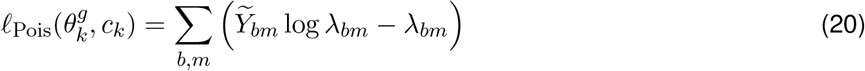

with respect to both 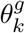 and *c*_*k*_, and then retaining the optimal 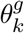. Here,

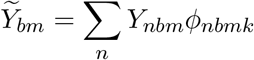

denotes the soft count assigned to process *k* in bin *b* and mutation type *m*, and

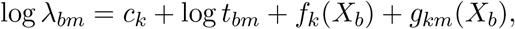

where *f*_*k*_ is treated as fixed throughout this update.

Defining

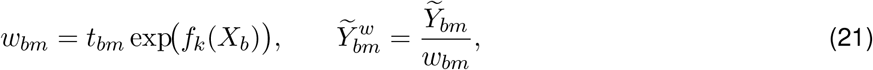

we can rewrite the objective as

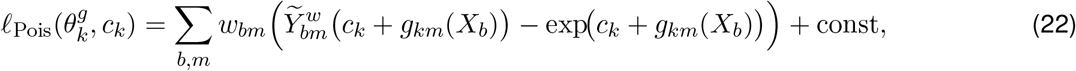

where const denotes terms independent of 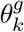 and *c*_*k*_. Since *g*_*km*_(*X*_*b*_) is linear in 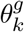 under the parameterization below, 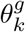 and *c*_*k*_ can be optimized jointly by weighted Poisson regression over bin–type pairs (*b, m*). Unlike for the non-linear macro-scale effects *f*_*k*_, it is not necessary to alternate between their optimizations or absorb the auxiliary scalar parameter *c*_*k*_ into the weights and responses.

#### Structure of the spectra effects

We decompose the spectra effects for process *k* into baseline, meso-scale, and strand-orientation terms,

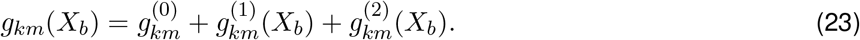

Here, 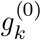 defines a baseline mutation spectrum for the process, 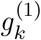 captures meso-scale genomic modulation, and 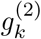 captures strand-dependent modulation.

To define these terms, let 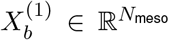 denote the meso-scale features and let 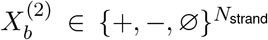 denote the strand-orientation features at bin *b*. For each mutation type *m* ∈ {1, …, 192}, let

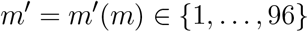

denote its *C/T* -centered representation, and let

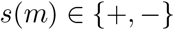

denote its strand configuration, with + for *C/T* -centered mutation types and − for *G/A*-centered mutation types.

#### Baseline spectrum

The baseline spectrum is parameterized by a vector 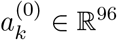, and is given by

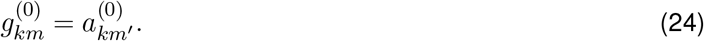

Thus, the baseline spectrum is shared between each mutation type and its *C/T* -centered reverse complement.

#### Meso-scale effects

For the meso-scale effects, we introduce an unpenalized shared coefficient vector

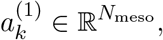

and an *ℓ*_1_-penalized mutation-type-specific coefficient matrix

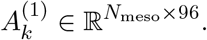

The resulting contribution is

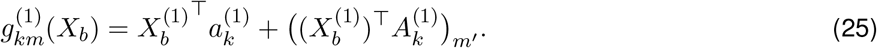

Thus, each meso-scale feature contributes both a mutation-type-independent shift and a mutation-type-specific deviation indexed by the *C/T* -centered mutation type.

#### Strand effects

For the strand effects, we introduce unpenalized shared coefficient vectors

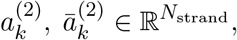

together with *ℓ*_1_-penalized mutation-type-specific coefficient matrices

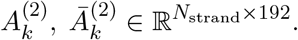

For each strand feature *i*, the contribution depends on whether the strand annotation at bin *b* matches the strand configuration of mutation type *m*. Writing 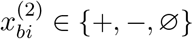 for the *i*-th strand feature, we define

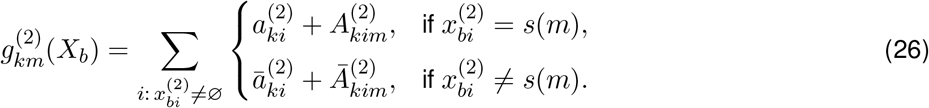

Unlike the meso-scale interaction coefficients, the strand-specific interaction terms 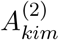 and 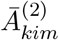 are indexed by the full mutation type *m* ∈ {1, …, 192}, since strand orientation distinguishes reverse-complement mutation types.

**Table 1** illustrates the coefficients chosen by a summand of (26) for a representative mutation type and its reverse complement: the two receive different coefficients depending on their orientation relative to the strand feature.

**Table 1:** Strand-dependent coefficients. Coefficients selected for a mutation type and its reverse-complement. The two receive different coefficients depending on whether the strand feature matches the mutation orientation.

| mutation type $m$ | $s(m)$ | strand feature $x_{bi}^{(2)}$ | | |
| --- | --- | --- | --- | --- |
| | | + | - | ∅ |
| $G[C > G]T$ | + | $a_{ki}^{(2)} + A_{kim}^{(2)}$ | $\bar{a}_{ki}^{(2)} + \bar{A}_{kim}^{(2)}$ | 0 |
| $A[G > C]C$ | - | $\bar{a}_{ki}^{(2)} + \bar{A}_{kim}^{(2)}$ | $a_{ki}^{(2)} + A_{kim}^{(2)}$ | 0 |

#### Optimization of the spectra effects *g*_*k*_

Under the parameterization above, log *λ*_*bm*_ is linear in the parameters

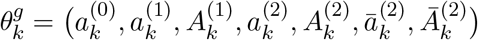

and *c*_*k*_. Therefore, for fixed *f*_*k*_, updating *g*_*k*_ is equivalent to solving a weighted Poisson generalized linear model with responses 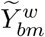, weights *w*_*bm*_, and linear predictor *c*_*k*_+*g*_*km*_(*X*_*b*_). We solve this problem using regularized iteratively reweighted least squares in the style of glmnet [6], applying *ℓ*_1_-penalization only to the mutation-type-specific interaction terms 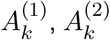, and 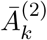, while leaving the shared coefficients and baseline spectrum unpenalized.

In summary, the spectra effects model combines an unpenalized baseline spectrum 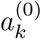, unpenalized shared feature effects 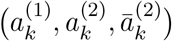, and sparse mutation-type-specific interaction terms 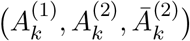. This design allows the model to capture broad systematic shifts in mutational spectrum while only introducing mutation-type-specific deviations from the baseline spectrum when strongly supported by the data.

### Scalable parameter estimation via stochastic variational inference

Algorithm 1 summarizes the overall coordinate-ascent structure of MuTopia model fitting, while Algorithm 2 makes the corresponding full-batch variational E-step and M-step updates explicit. In the full-batch procedure, all variational and model parameters are updated using the complete dataset at every iteration. Although these updates monotonically increase the ELBO, this becomes computationally expensive for large datasets.

To improve scalability, we therefore use stochastic variational inference (SVI; Algorithm 3), following the general framework of Hoffman et al. [7]. At each iteration, we subsample a set of genomic bins and restrict the mutation counts and features to those bins. We then apply the same variational E-step and M-step structure as in the full-batch algorithm, but on the subsampled data. To obtain an unbiased estimate of the corresponding full-data update, the minibatch contribution to the variational parameter *β*_*n*_ is rescaled by the inverse subsampling rate. The resulting stochastic parameter estimates are then merged with the current global parameters using the learning rate *ρ*_*i*_ = (*i* + 1)^−0.5^. Under standard conditions for stochastic approximation, these updates converge to a local optimum of the variational objective provided that they yield unbiased estimates of the natural gradient [7]. In our setting, this condition is satisfied by bin subsampling together with the rescaling step above. After convergence, sample-specific variational parameters can be obtained by a final E-step on the full data using the fitted global parameters. In practice, SVI substantially reduces training time and often yields better solutions than the full-batch procedure.

#### Algorithm 1

MuTopia — coordinate ascent overview.

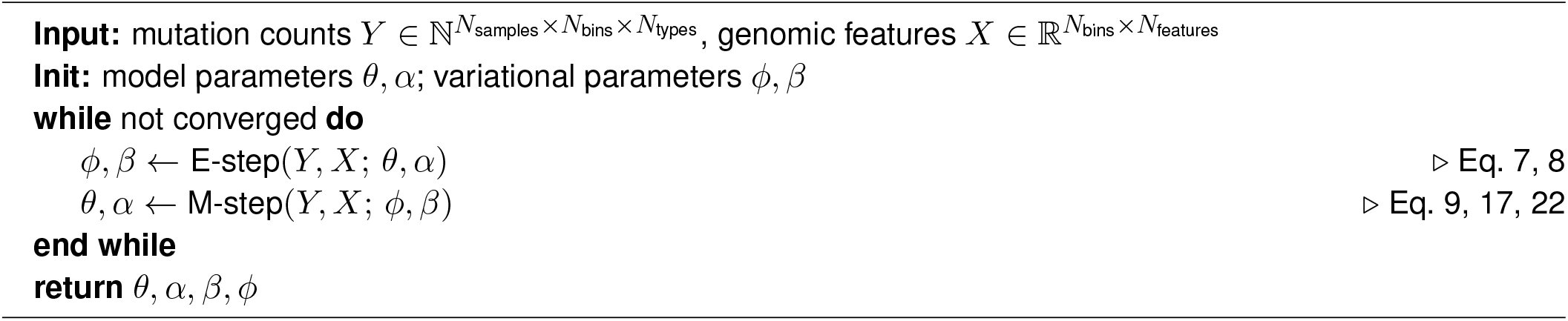

#### Algorithm 2

MuTopia — full-batch coordinate ascent variational inference.

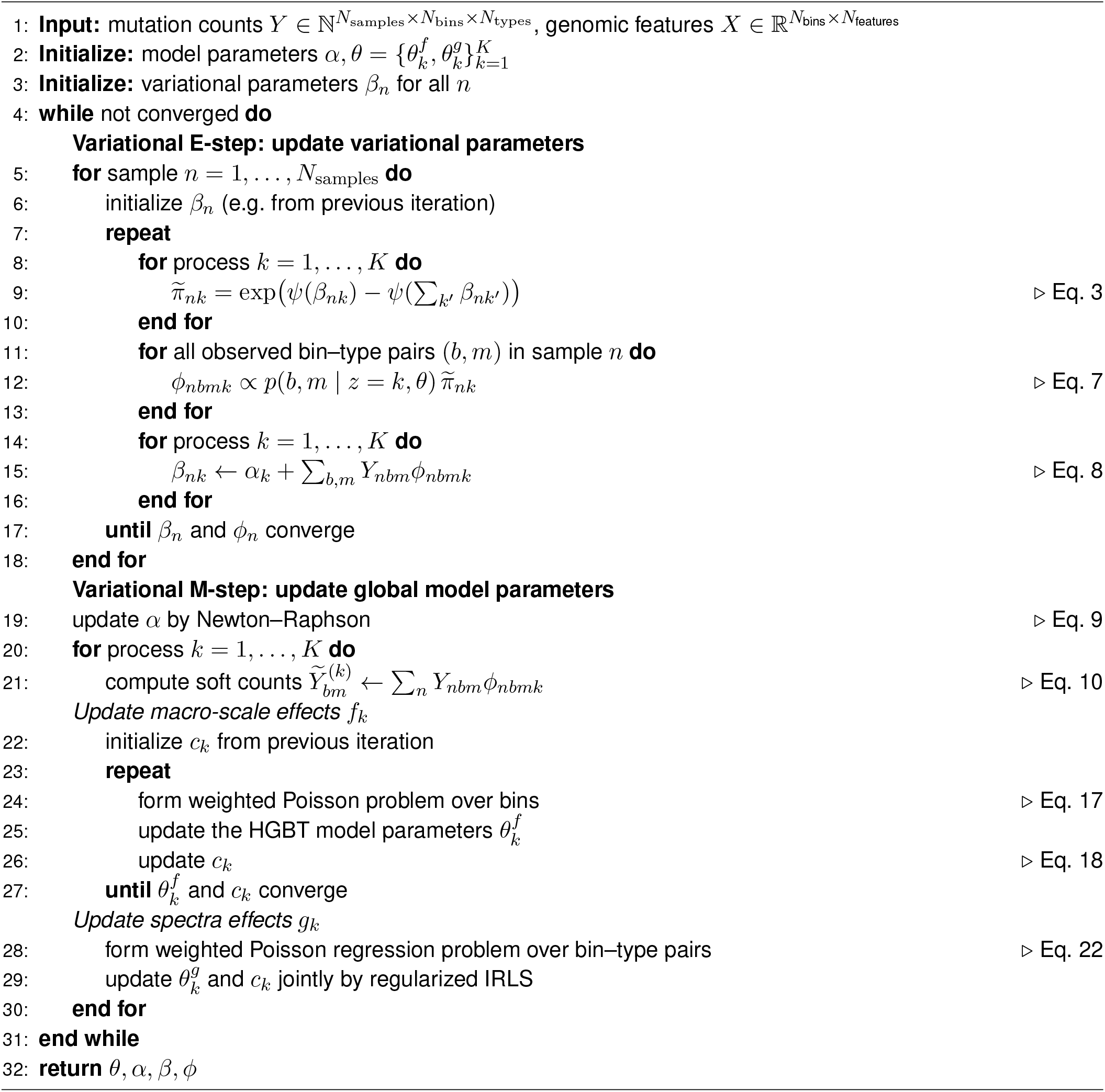

#### Algorithm 3

MuTopia — stochastic variational inference.

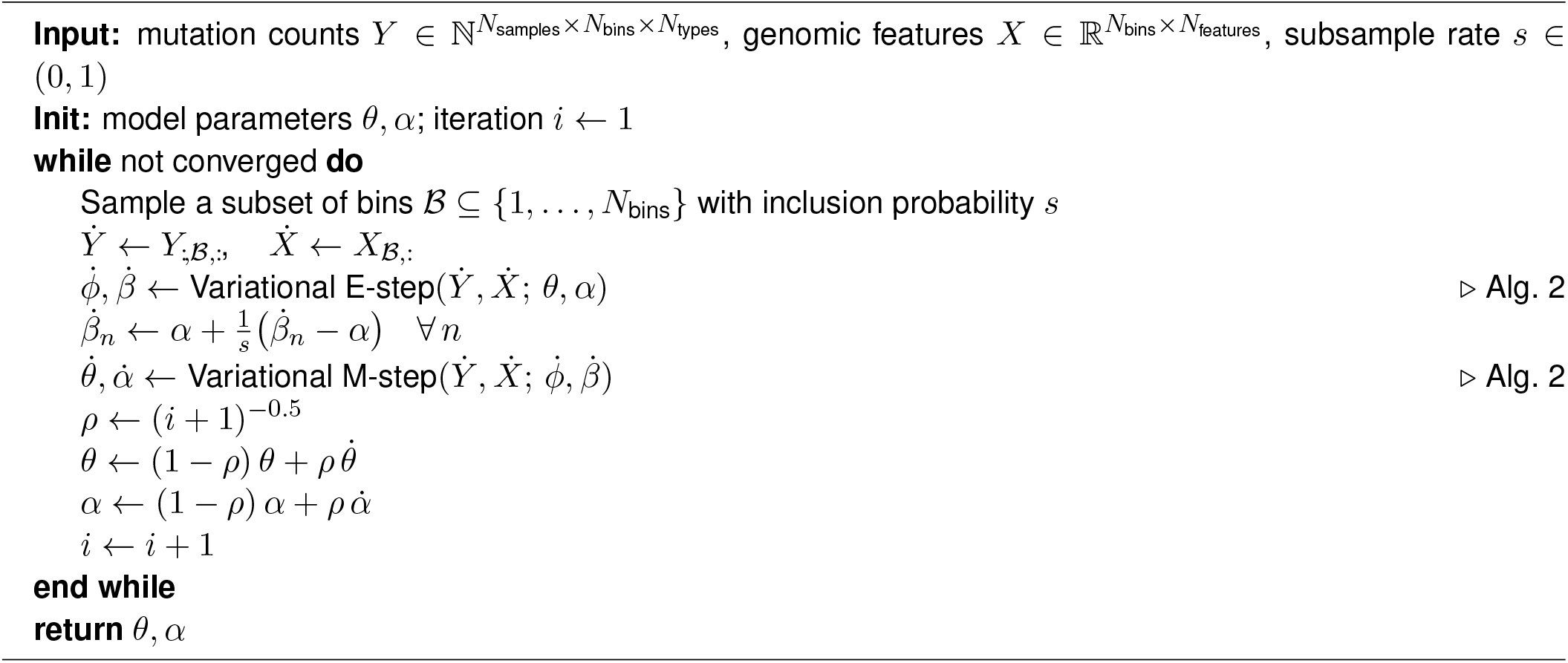

### Feature selection from ENCODE

We implemented a fully automated strategy to build tissue-matched corpora of genetic and epigenetic features for each dataset used in our study, following the AlphaGenome [8] approach of prioritizing samples based on their ENCODE-assessed quality control metrics. Briefly, we fetched unperturbed ENCODE experiments for DNase-seq, ATAC-seq, Whole-genome Bisulfite Sequencing (WGBS), Histone ChIP-seq, Repli-seq, Poly-A plus RNA-seq, and total RNA-seq assays. Using the ENCODE audit metadata, we filtered out experiments that failed to meet QC, read length, or fraction of reads in peaks (FRiP) thresholds, as outlined in AlphaGenome. We then assigned each experiment a quality score between −1 (FAIL) and 4 (PASS). Next, we grouped the experiments by biosample ontology term and assay. For each group, we chose the single most concordant experiment that best satisfied the following criteria:

1. Oldest life stage (“adult” prioritized over “child”)
2. The description contained none of the terms “genetically modified”, “arrested”, or “treated”.
3. Maximized quality score.
4. Used paired-end reads.
5. Conducted in tissue or primary cells.
6. Did not measure a subcellular fraction.

For experiments that met those criteria, we then selected the one with the highest FRiP, if applicable, or the most recently conducted.

Finally, for each tumor type in the PCAWG cohort, we specified a ranked list of Uberon tissue ontology terms [9] (**Table 2**). To choose the best-matched experiment to represent a tumor cell type, we used the following prioritization heuristic, selecting the experiment that maximally satisfied the conditions:

**Table 2:** Uberon cell type ontology terms — used to query ENCODE experiments for each tumor type.

| TumorType | Tissue terms (ranked) | Repli-seq terms |
| --- | --- | --- |
| Bladder-TCC | UBERON:0001255; UBERON:0001259;<br>UBERON:0005033; CL:2000040 | EFO:0001187 |
| Cervix-All | UBERON:0000995; EFO:0002791 | EFO:0002791 |
| ColoRect-AdenoCA | UBERON:0000317; UBERON:0004992;<br>UBERON:0001159; UBERON:0001157;<br>UBERON:0008971 | EFO:0001187 |
| Eso-AdenoCA | UBERON:0001043; UBERON:0002469;<br>UBERON:0004648 | EFO:0001187 |
| Head-SCC | UBERON:0006920 | EFO:0001196 |
| Breast-All | UBERON:0008367; UBERON:0000310;<br>CL:0002327 | EFO:0001203 |
| Kidney-All | UBERON:0002113; UBERON:0004538;<br>UBERON:0004539; UBERON:0001225;<br>CL:0002518; CL:1000510; CL:1000892;<br>UBERON:0001255 | EFO:0001187 |
| Liver-HCC | UBERON:0002107; UBERON:0001115;<br>UBERON:0001114 | EFO:0001187 |
| Lung-All | UBERON:0002048; UBERON:0002168;<br>UBERON:0002167; UBERON:0008953;<br>UBERON:0008952; UBERON:0002170;<br>UBERON:0002171 | EFO:0001196 |
| Ovary-AdenoCA | UBERON:0000992 | EFO:0001203 |
| Panc-AdenoCA | UBERON:0001264; UBERON:0001150 | EFO:0001187 |
| Prost-AdenoCA | UBERON:0002367; CL:0002231 | EFO:0001203 |
| Skin-Melanoma | CL:1000458; UBERON:0002097;<br>UBERON:0001003; UBERON:0004264;<br>UBERON:0036149 | EFO:0001196 |
| Stomach-AdenoCA | UBERON:0000945 | EFO:0001187 |
| Uterus-AdenoCA | UBERON:0000995; EFO:0002791 | EFO:0002791 |

1. Life stage was “adult”.
2. Quality score exceeded 3.
3. Highest rank of tissue ontology term.
4. Highest quality score.

For replication timing data, we utilized the best-matched cell line in which each phase had been assayed. We used the *LiftOver* [10] tool to convert the hg19-aligned Repli-seq “percentage normalized signal” files to hg38 coordinates in line with the rest of our analysis.

All together, **Table 3** summarizes how these genomic features were used in the MuTopia model. Macro features (continuous-valued) capture megabase-scale variation in mutation rate. Meso and strand features modulate the mutation type distribution within a bin.

**Table 3:** Genomic feature overview. Features used in the MuTopia model, grouped by feature type (macro, meso, strand). Log1p-CPTM: log1p-counts per ten million; quantile: quantile normalization; Repeat element fraction is the fraction of soft-masked bases in the reference genome for a given bin. ^*†*^Replication strand was derived from MCF7 Repli-seq and is not tissue-matched for non-breast tumor types.

| Feature | Biological signal | Feature type | Tissue-matched | Assay/Source | Normalization | Affects binning | Domain |
| --- | --- | --- | --- | --- | --- | --- | --- |
| Gene expression | Transcriptional activity | Macro | Yes | RNA-seq | Log1p-CPTM | No | continuous |
| Chromatin activity | Diffuse chromatin accessibility | Macro | Yes | DNase-seq | Log1p-CPTM | No | continuous |
| Repli-seq (G1b, S1-S4, G2) | Replication timing phases | Macro | Yes | Repli-seq | quantile | No | continuous |
| H3K27ac, H3K4me1, H3K4me3 | Active enhancers and promoters | Macro | Yes | ChIP-seq | Log1p-CPTM | No | continuous |
| H3K36me3, H3K27me3, H3K9me3 | Heterochromatin marks | Macro | Yes | ChIP-seq | Log1p-CPTM | No | continuous |
| CpG methylation fraction | DNA methylation | Macro | Yes | WGBS | standardized | No | continuous |
| GC content | Nucleotide composition | Macro | No | Reference genome | standardized | No | continuous |
| Repeat element fraction | Repeat element density | Macro | No | Reference genome | standardized | No | continuous |
| ATAC accessibility | Focal open chromatin regions | Meso | Yes | ATAC-seq | quantile | Yes | binary |
| Gene strand | Transcription orientation | Strand | No | MANE annotation [11] | — | Yes | {+, -, ∅} |
| Replication strand | Replication fork direction | Strand | No<sup>†</sup> | Repli-seq (MCF7) | — | Yes | {+, -, ∅} |

### Software and implementation

MuTopia was written in Python, with numerical operations implemented using *numpy, scipy, sparse* and *numba* [12–14]. Hyperparameter optimization was managed using *optuna* [15]. Data organization was facilitated using *xArray, netCDF4*, and *Pandas* [16–18]. Dataset configuration and construction were implemented using *pydantic, pyyaml*, and *luigi*. Plotting and analyses were conducted with *matplotlib, jupyter*, and *pygenometracks* [19, 20].

## Supplementary figures

**Supplementary Figure 1:**
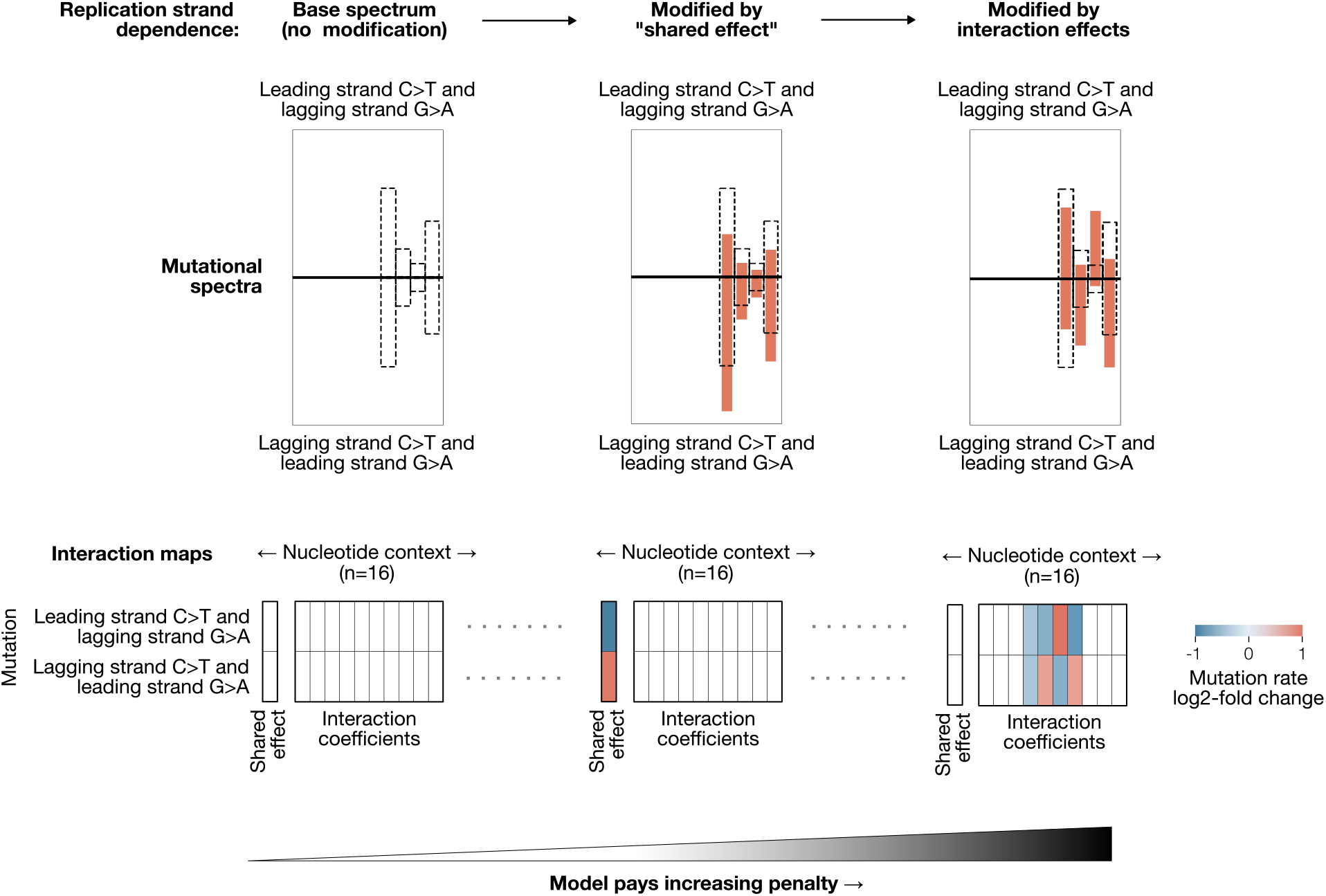
Strand effect parameterization and regularization strategy. MuTopia learns context-dependent signatures in which mutation rates are modified by local genomic state on a per-mutation-type basis. To reduce the variance of the model when fitting these complex signatures, we introduce a hierarchical method for learning sparse “interaction maps”. First, no context dependence incurs no regularizing penalty (left). Then, uniform scaling across all mutation types (middle) is favored over the fitting of type-specific interactions (right).

**Supplementary Figure 2:**
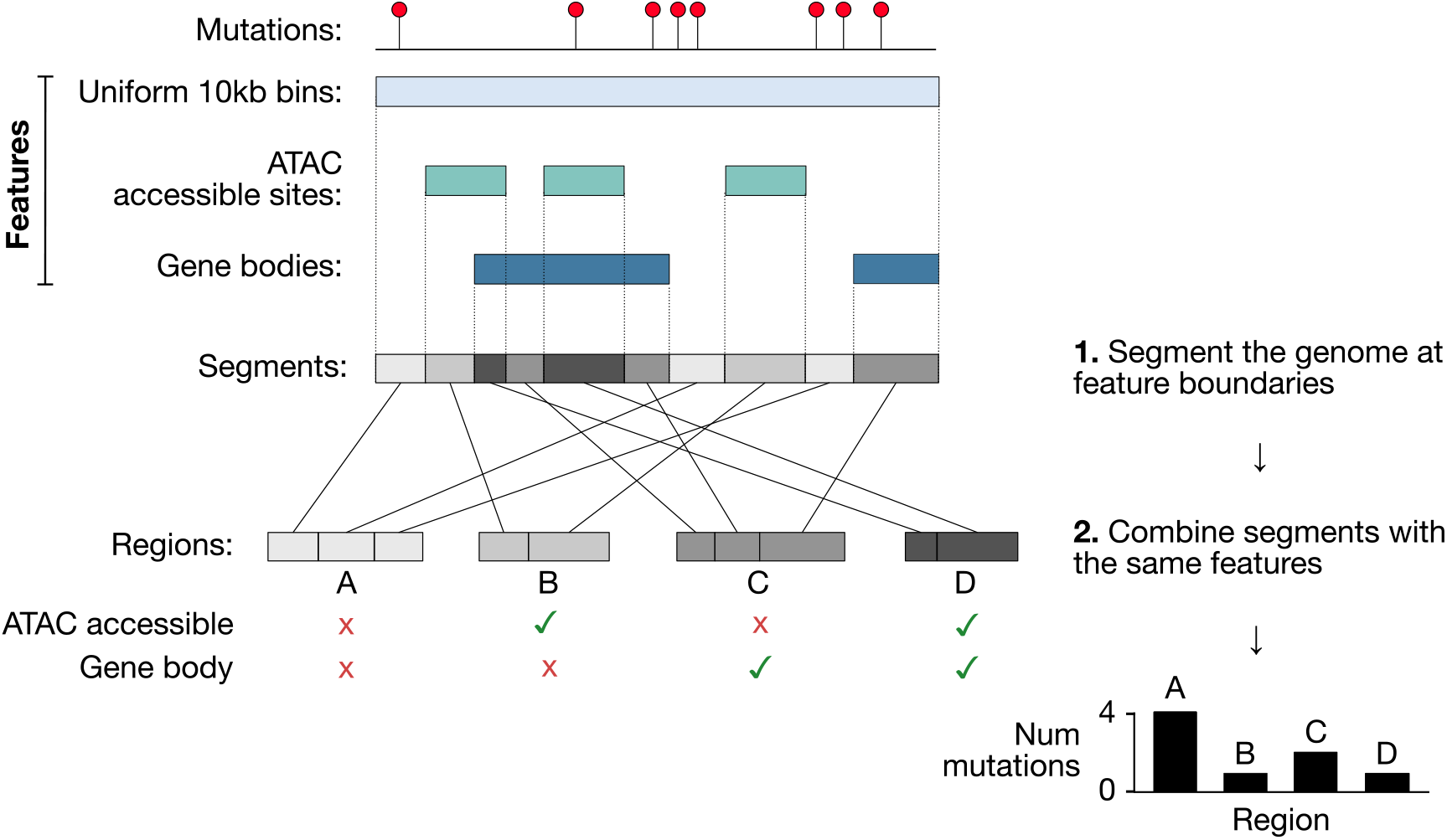
MuTopia genome-binning and mutation aggregation strategy. For tractable parameter updates, we locally aggregate mutation counts into discontinuous bins that share the same locus features. The resolution of the model (in this case 10kb) determines the maximum distance at which any two mutations may be aggregated. The macro-scale bins are segmented according to the intersection of any underlying discrete genomic features. Then, mutations within segments which intersect the same sets of features are pooled. The segment groups are indicated by color.

**Supplementary Figure 3:**
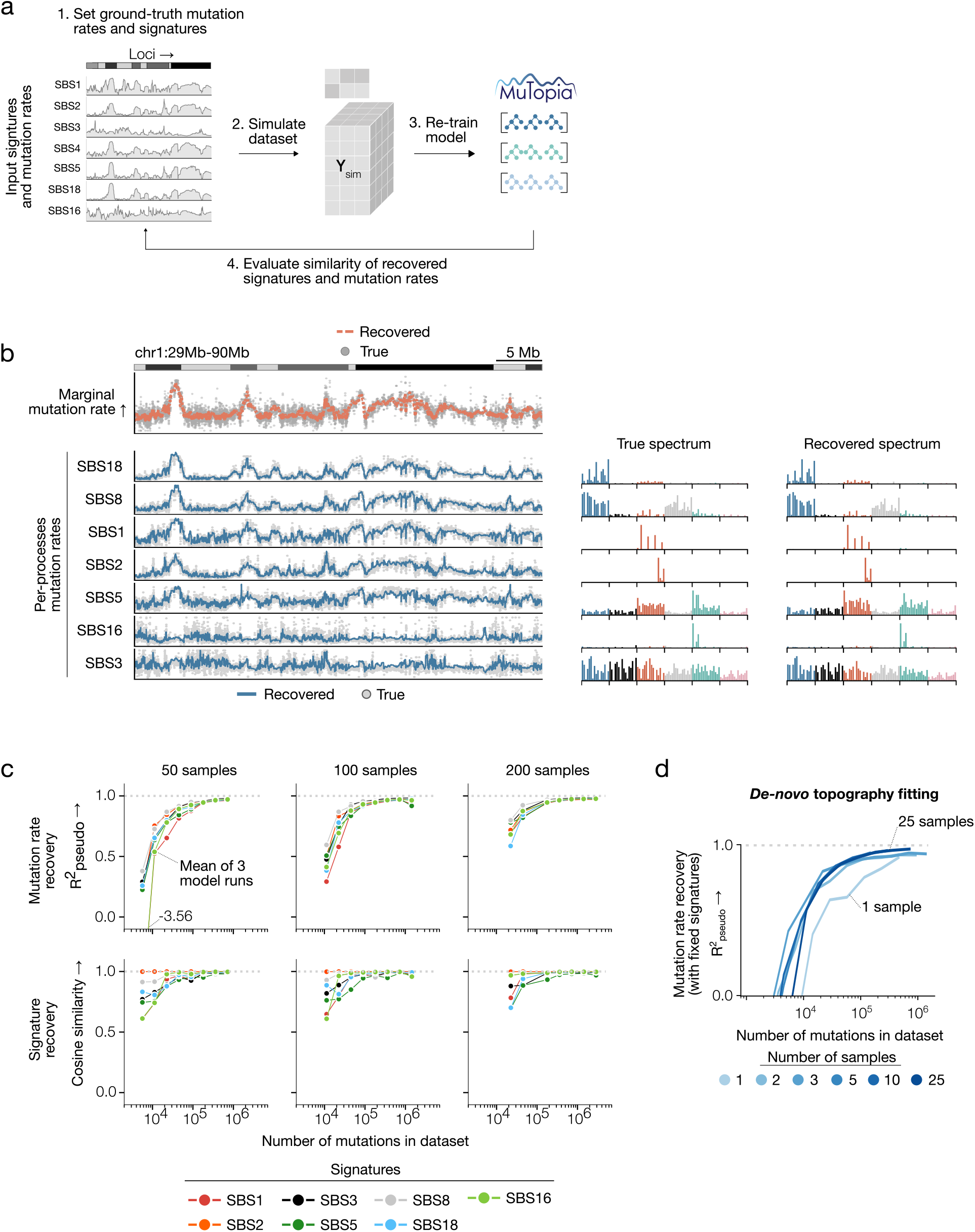
Simulation-based benchmarking results. **a)** Schematic of the simulation-based evaluation strategy. Using realistic components fit from real breast cancer data, we sampled new datasets while varying the process contributions and burdens. We trained new models on these simulated datasets, then compared the similarity between the extracted components and those input into the simulation using pseudo-*R*^2^ between mutation rate distributions and cosine similarity between spectra. **b)** The recovered mutational spectra and mutation rate distributions were compared to input data in unseen regions of the genome. **c)** Mutation rate and spectrum recovery quality, broken down by dataset size and by input signature for varying numbers of simulated samples. **d)** Performance on *de novo* topography fitting while fixing signatures.

**Supplementary Figure 4:**
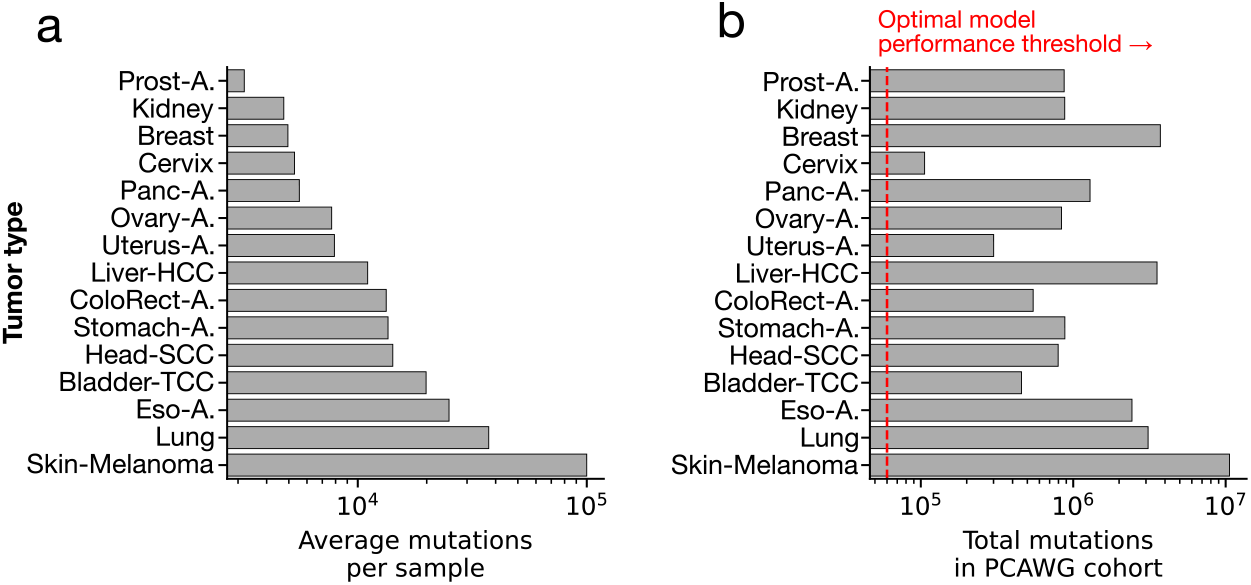
Pan-cancer dataset properties. **a)** Average number of mutations per sample in the dataset. **b)** Total number of mutations per tumor type.

**Supplementary Figure 5:**
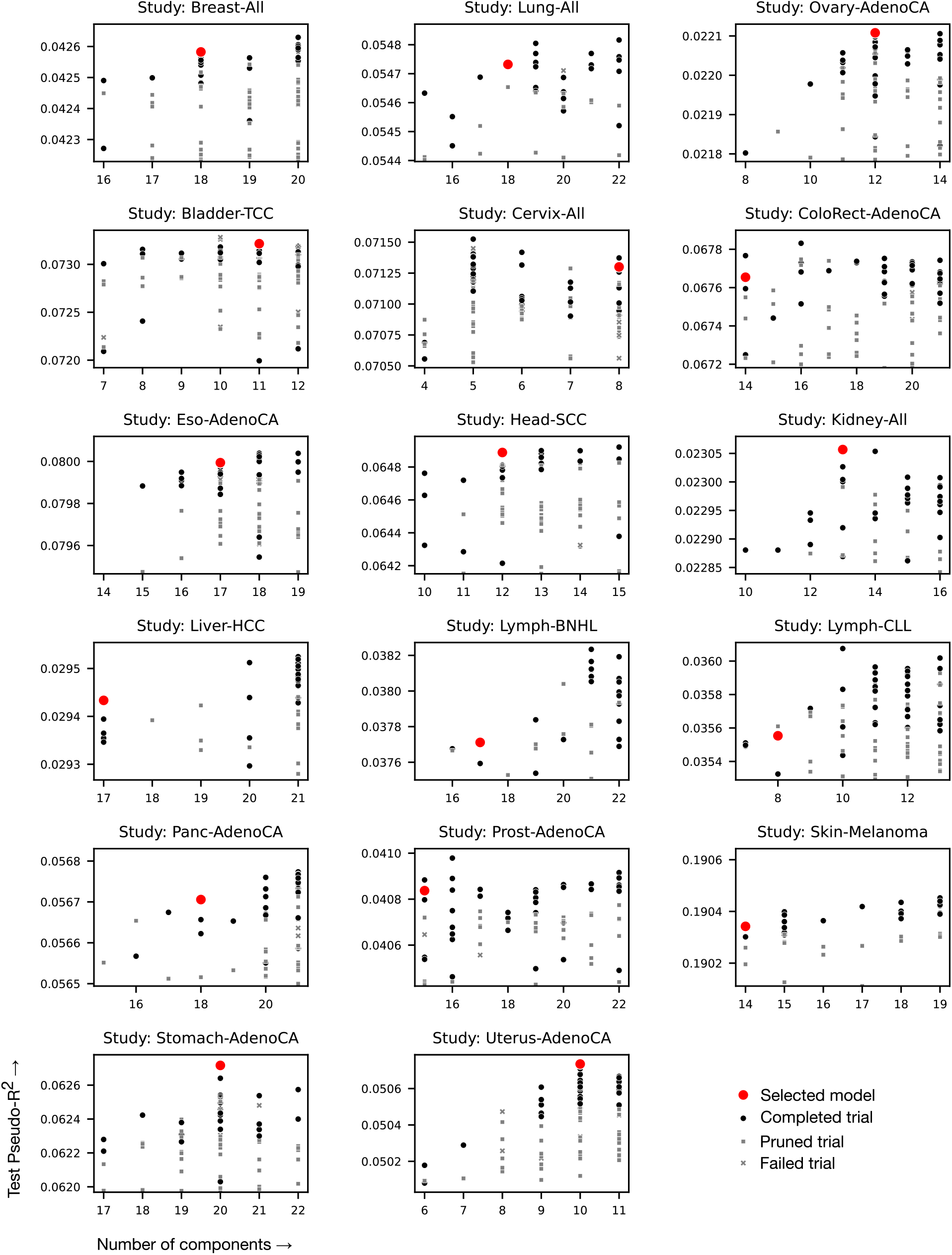
Pan-cancer hyperparameter optimization results. Number of components versus score on held-out chromosome 2 for the hyperparameter optimization “studies” performed on each tumor type.

**Supplementary Figure 6:**
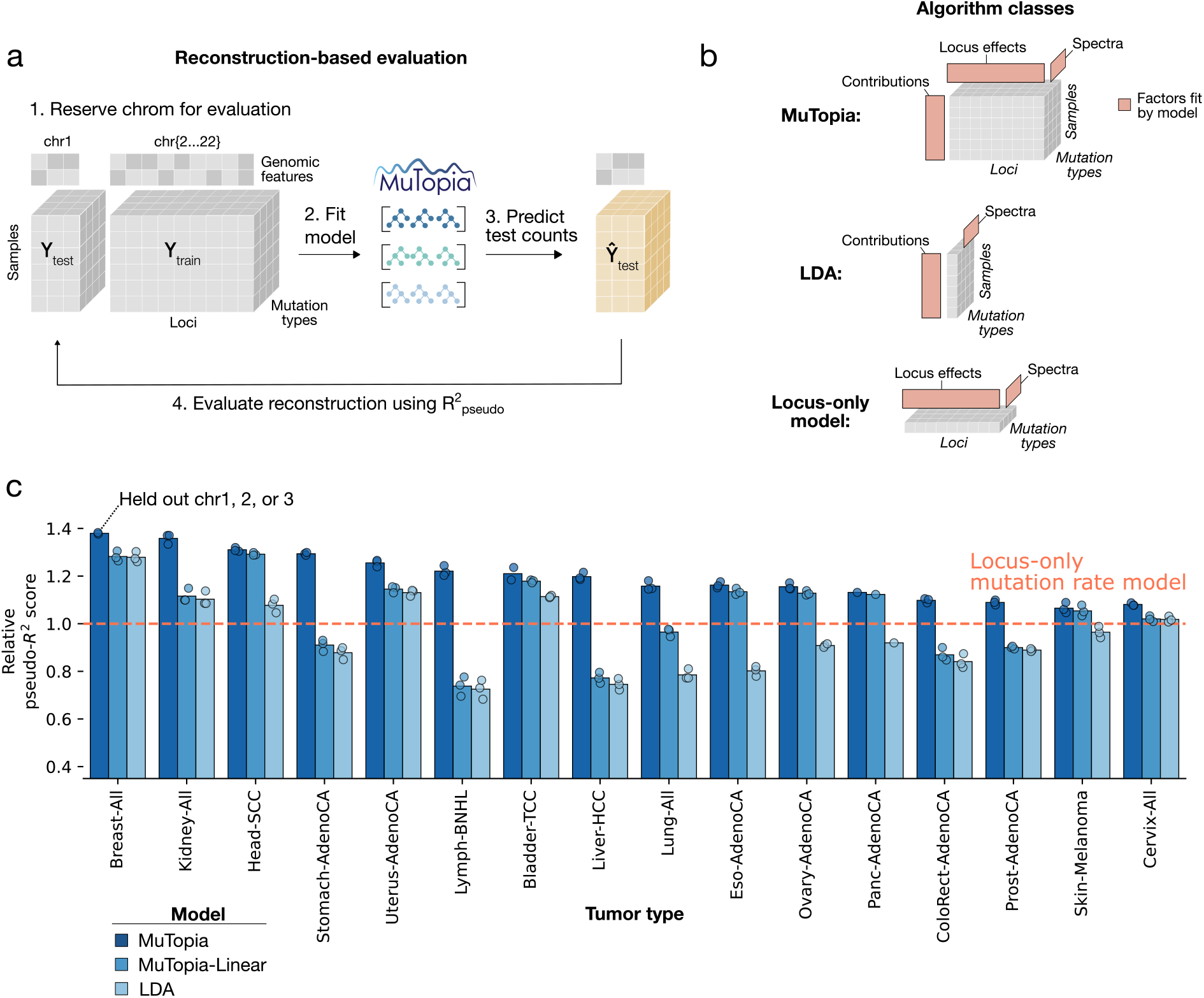
Reconstruction-based evaluation and model ablation results. **a)** Reconstruction-based evaluation method. Test data were reserved by chromosome, then imputed from a trained model. **b)** Algorithm classes evaluated in benchmarking tests. MuTopia fits three factors: process contributions, locus effects, and spectra. LDA/NMF omits locus effects, while “locus only” omits sample-level contributions. **c)** Results from reconstruction-based evaluation using the relative pseudo-*R*^2^ metric. Bars show the full and ablated versions of the MuTopia model. The dashed line shows the likelihood of the data using a baseline locus-based mutation rate model.

**Supplementary Figure 7:**
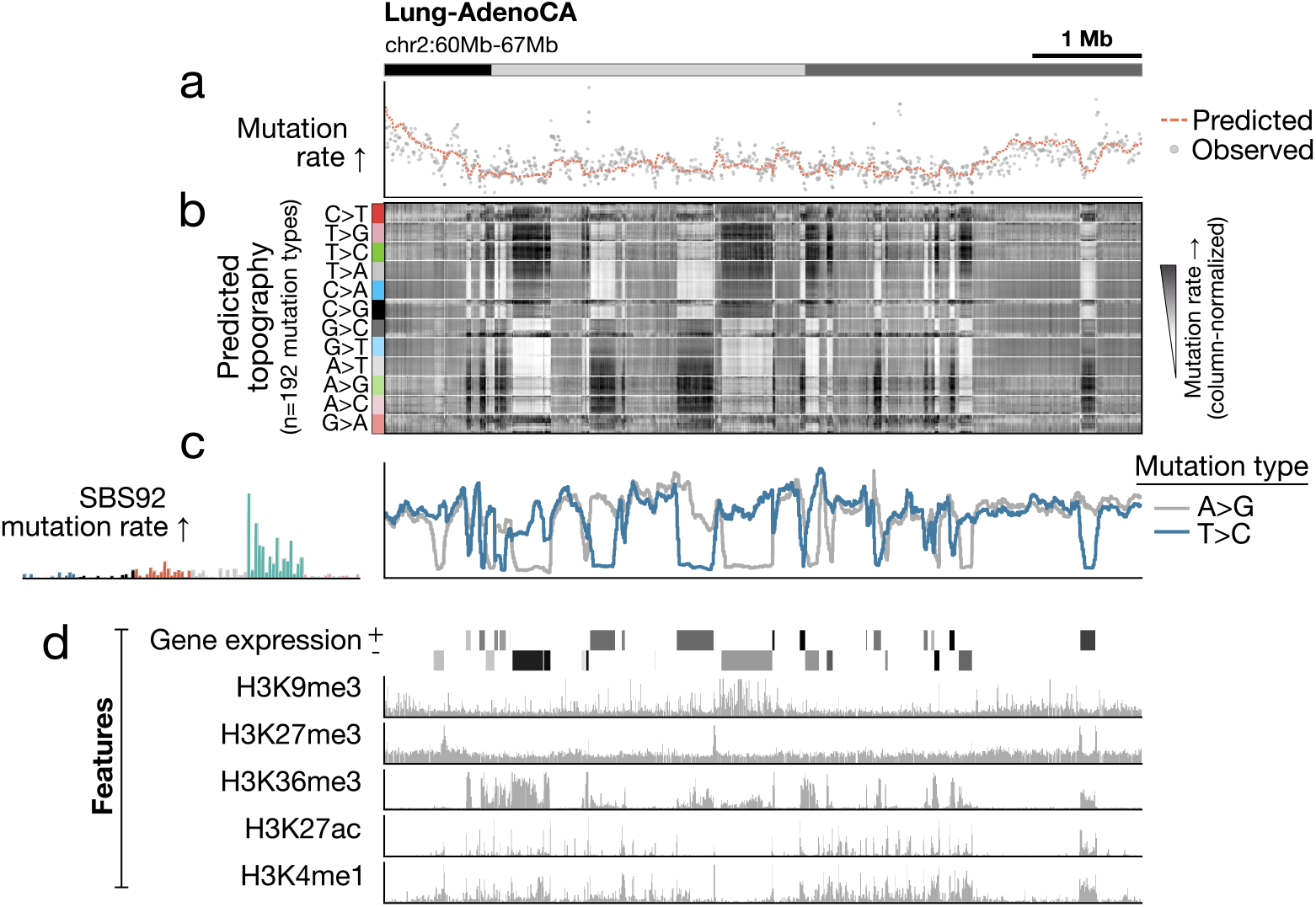
Mutational topography at the intermediate scale. **a)** Observed versus predicted mutation rates on a held-out section of the genome. **b)** Mutational topography prediction. **c)** Predicted mutation rate profiles of A*>*G versus T*>*C mutations originating from SBS92. **d)** Representative genomic features driving variation in mutation rates at this scale.

**Supplementary Figure 8:**
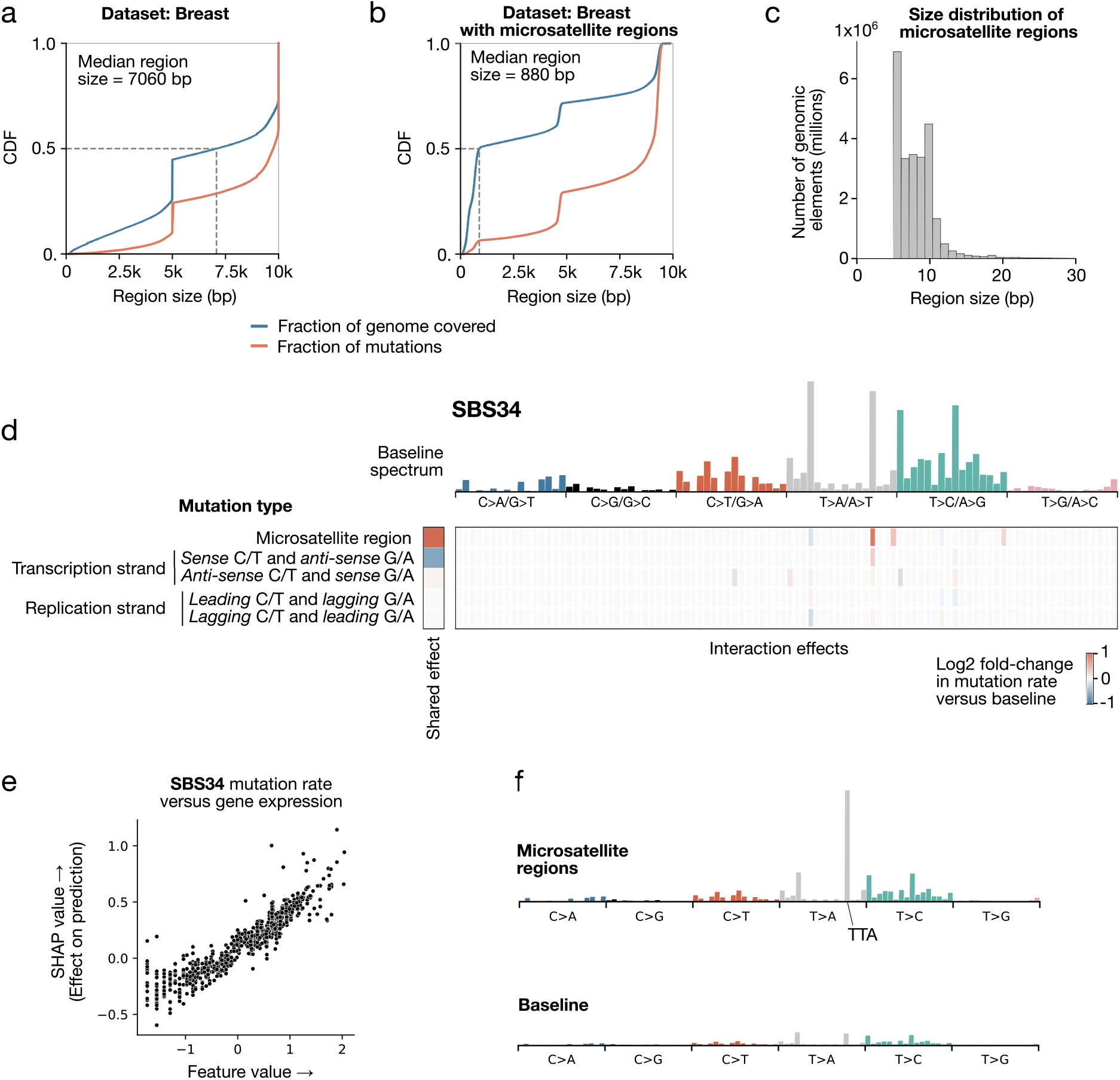
*Meso*-scale analysis of mutagenesis at microsatellite sites. **a)** For the breast dataset, cumulative distribution functions of fraction of genome covered by segmented regions versus the fraction of mutations those bins contain, sorted by region size. **b)** Same as (a), except with the addition of *meso*-scale features indicating microsatellite regions. **c)** Size distribution of microsatellite regions. **d)** Context interaction map for signature SBS34. **e)** Gene expression Shapley value versus feature value for signature SBS34. **f)** SBS34 signature mutational spectra in a “baseline” genomic context (bottom), and in microsatellite regions (top).

**Supplementary Figure 9:**
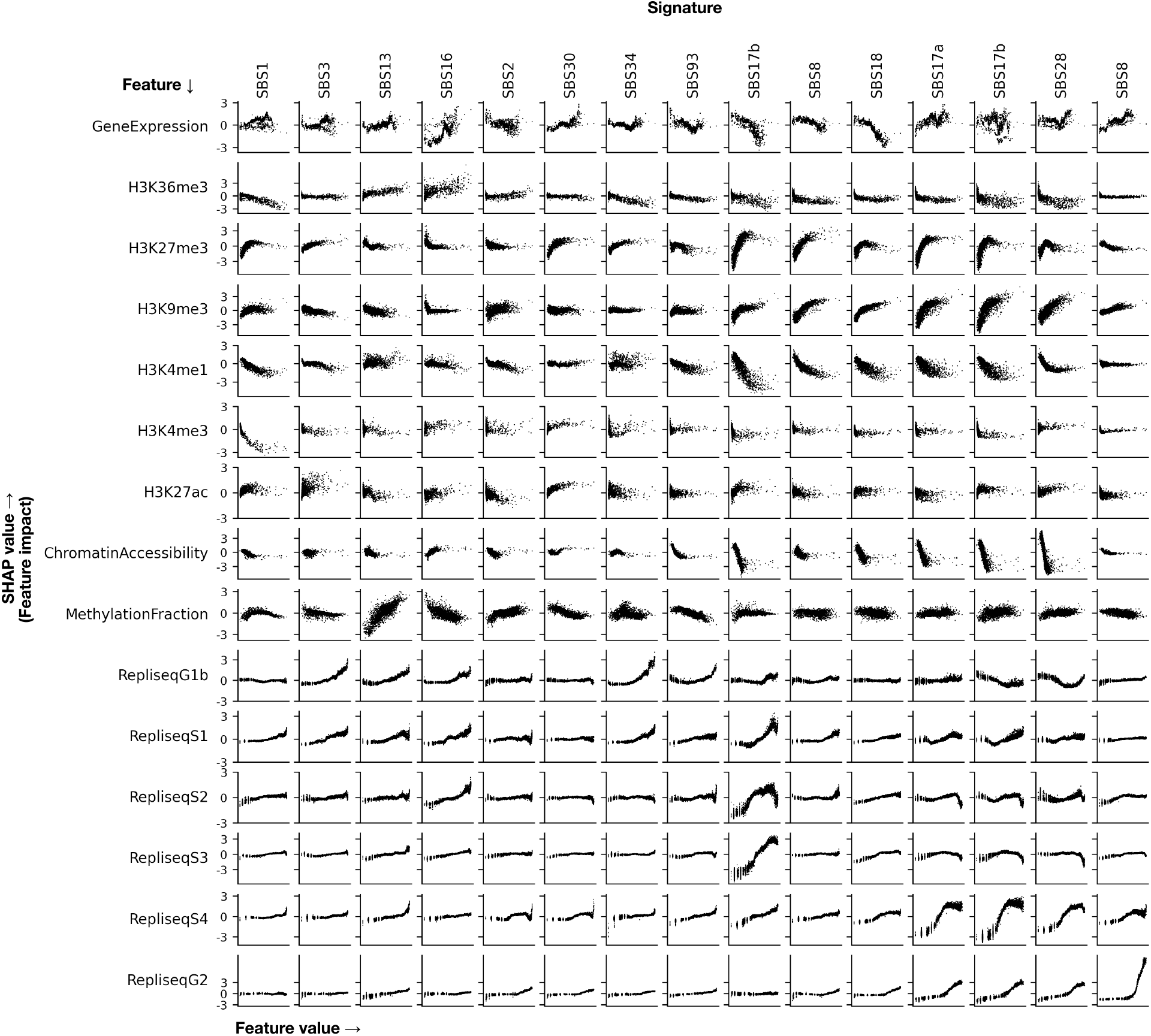
Stomach adenocarcinoma Shapley value analysis per process, per genomic feature. Shapley values inferred for each signature and feature combination in the stomach adenocarcinoma dataset. Each point is a genomic region, which is assigned a feature value and feature impact for each signature.

**Supplementary Figure 10:**
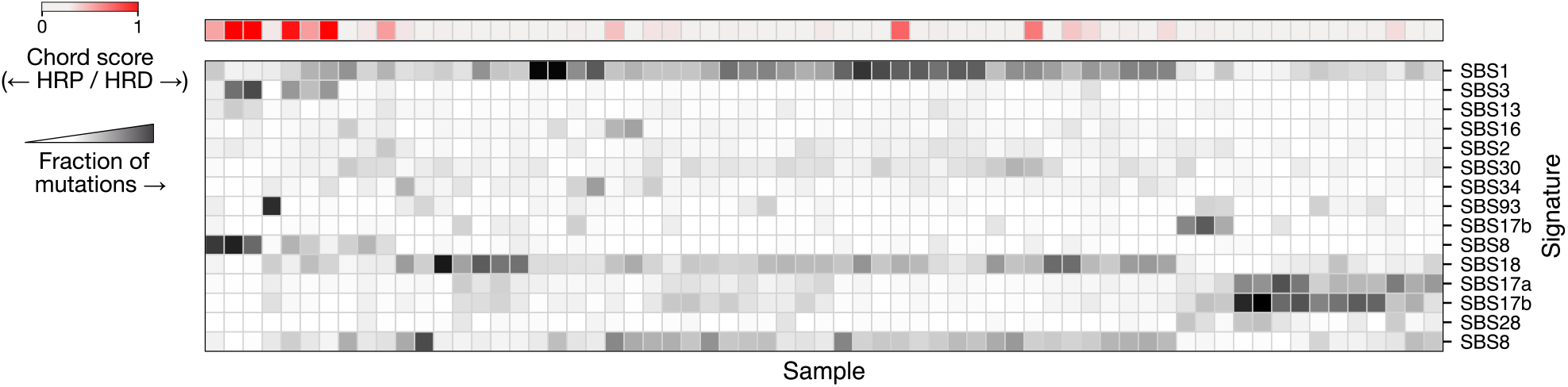
Relative contributions per process in stomach adenocarcinoma samples. (top) CHORD HRD likelihood for each stomach adenocarcinoma sample. (bottom) Fraction of mutations assigned to each component.

**Supplementary Figure 11:**
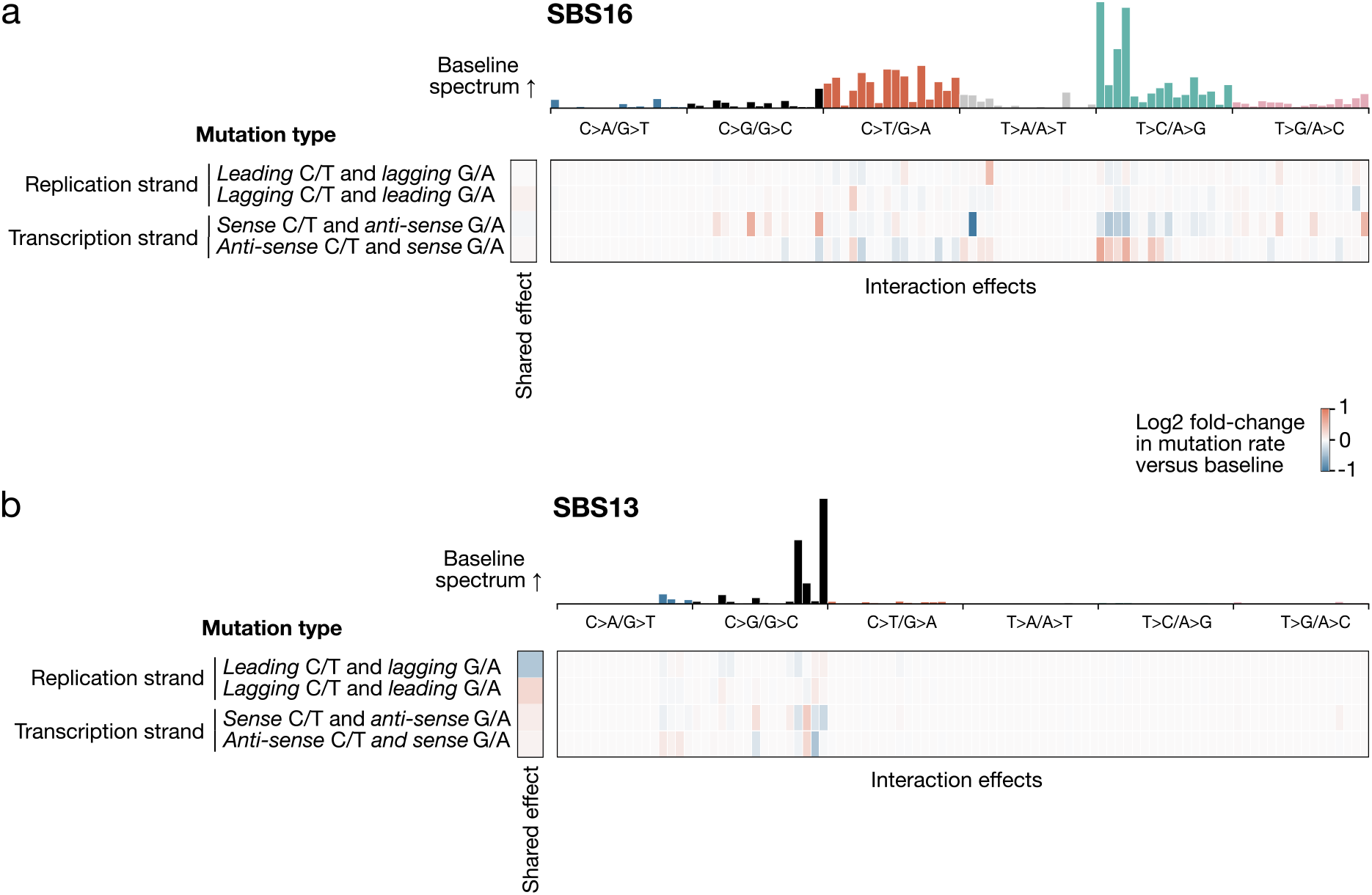
Context interaction maps. Context interaction maps for signatures SBS16 (**a**) and SBS13 (**b**) from the stomach adenocarcinoma dataset.

**Supplementary Figure 12:**
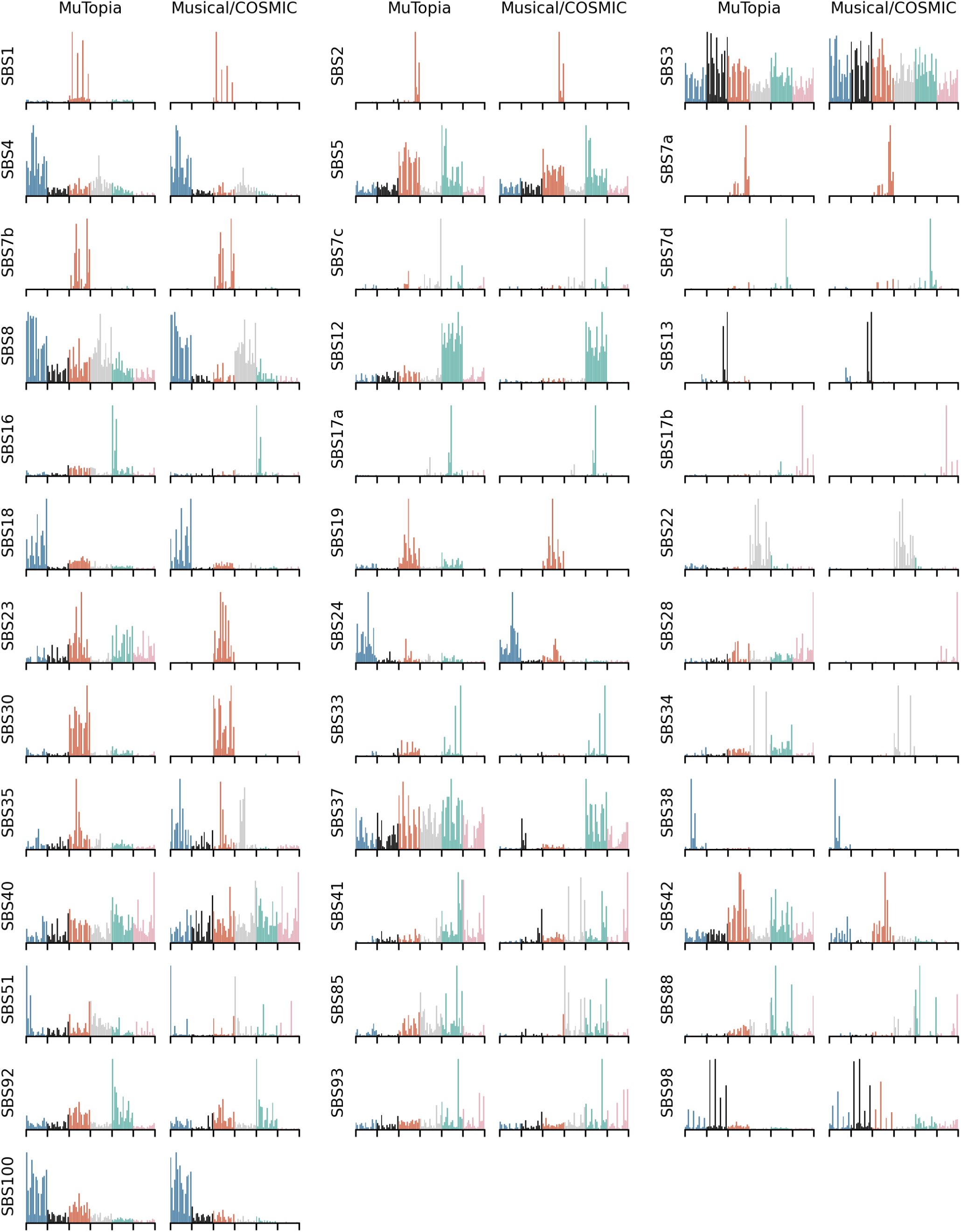
COSMIC-matching signature spectra. The average mutational spectrum across all MuTopia components assigned to a COSMIC signature versus the MuSiCal/COSMIC representation.

**Supplementary Figure 13:**
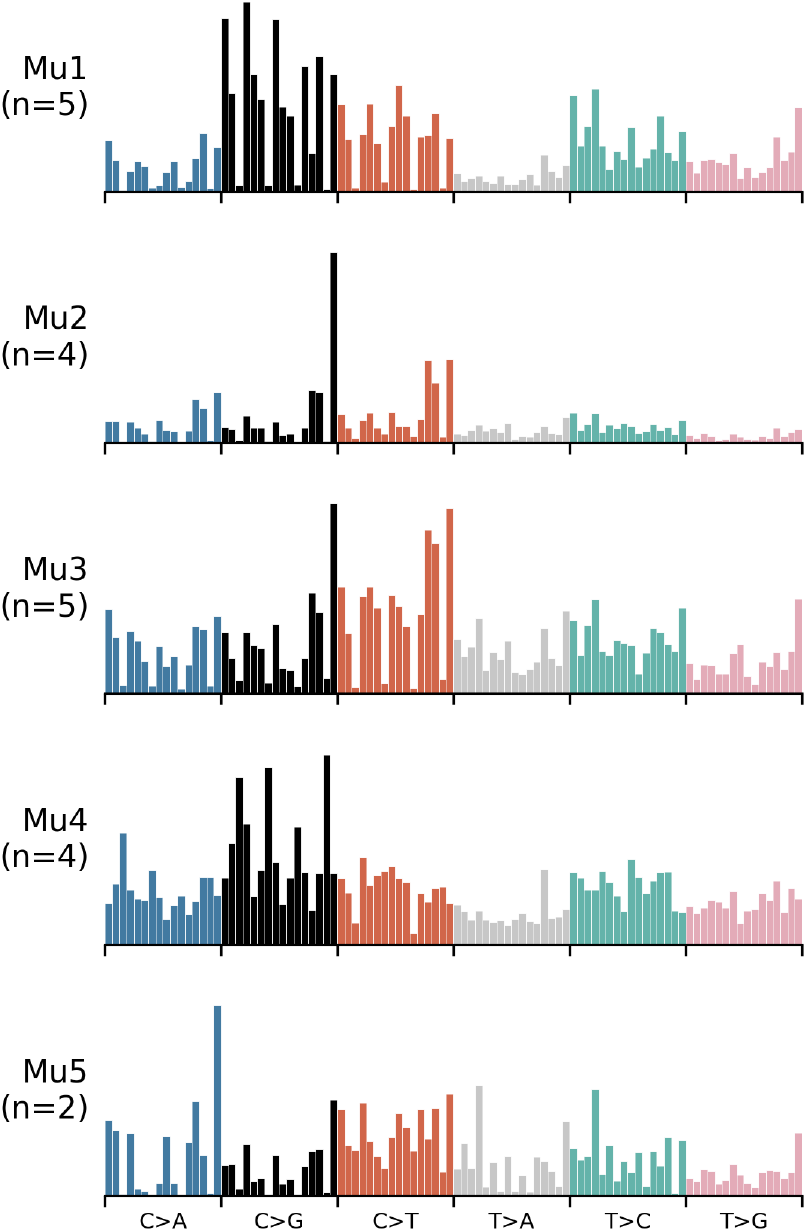
Novel mutational signature spectra found in pancancer analysis. Novel mutational signatures recurrently discovered by MuTopia in multi-cancer meta-analysis, where “n” denotes the number of tumor types in which each process was found.

**Supplementary Figure 14:**
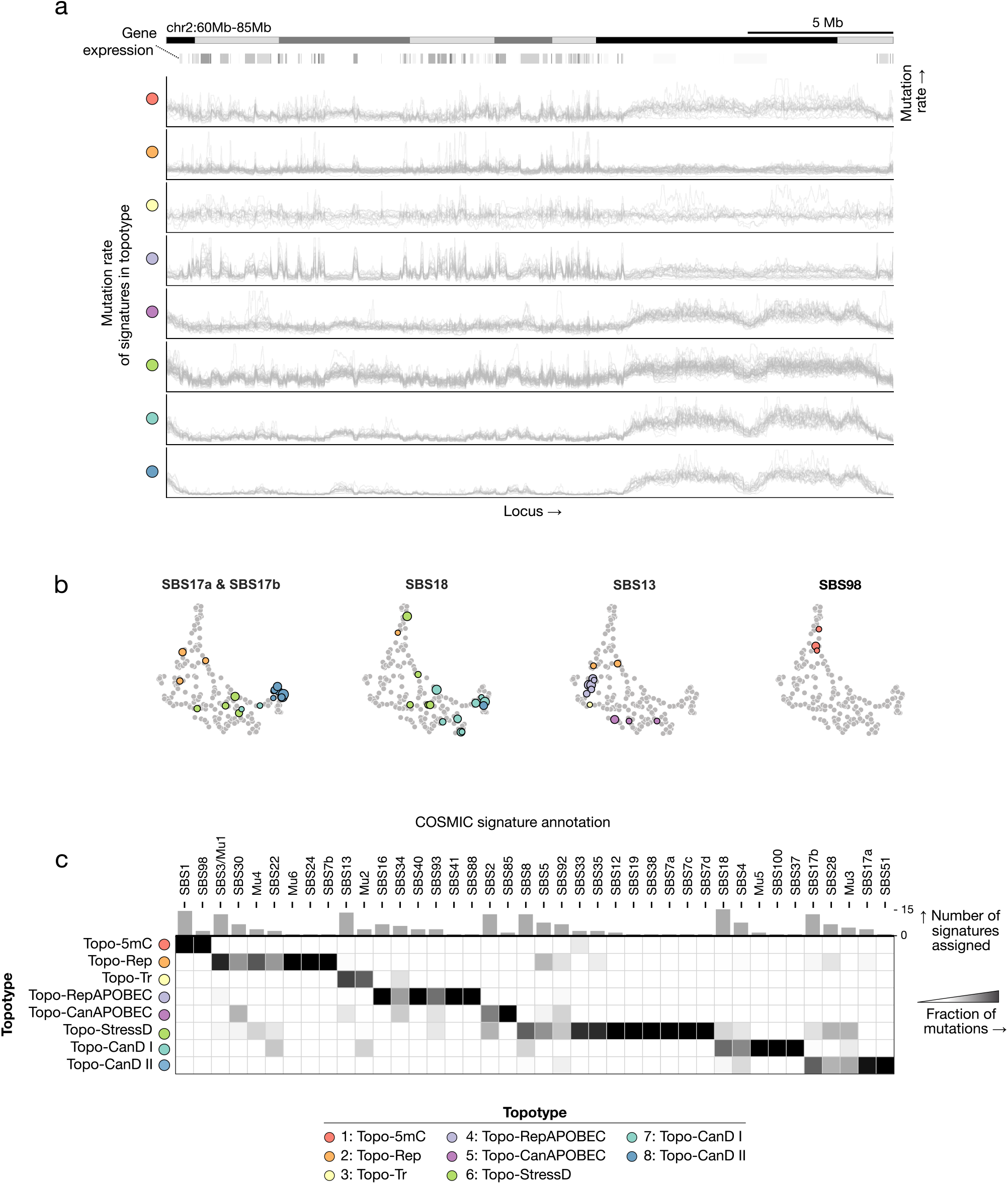
Signature topotype assignments. **a)** Mutation rate profile for each signature for each topotype. **b)** Fraction of mutations attributed to signatures of each topotype, grouped by those signatures’ COSMIC annotation. **c)** Signatures corresponding to COSMIC SBS annotations overlaid on the UMAP. For the points representing the highlighted signatures, the marker size indicates the log-transformed number of mutations.

**Supplementary Figure 15:**
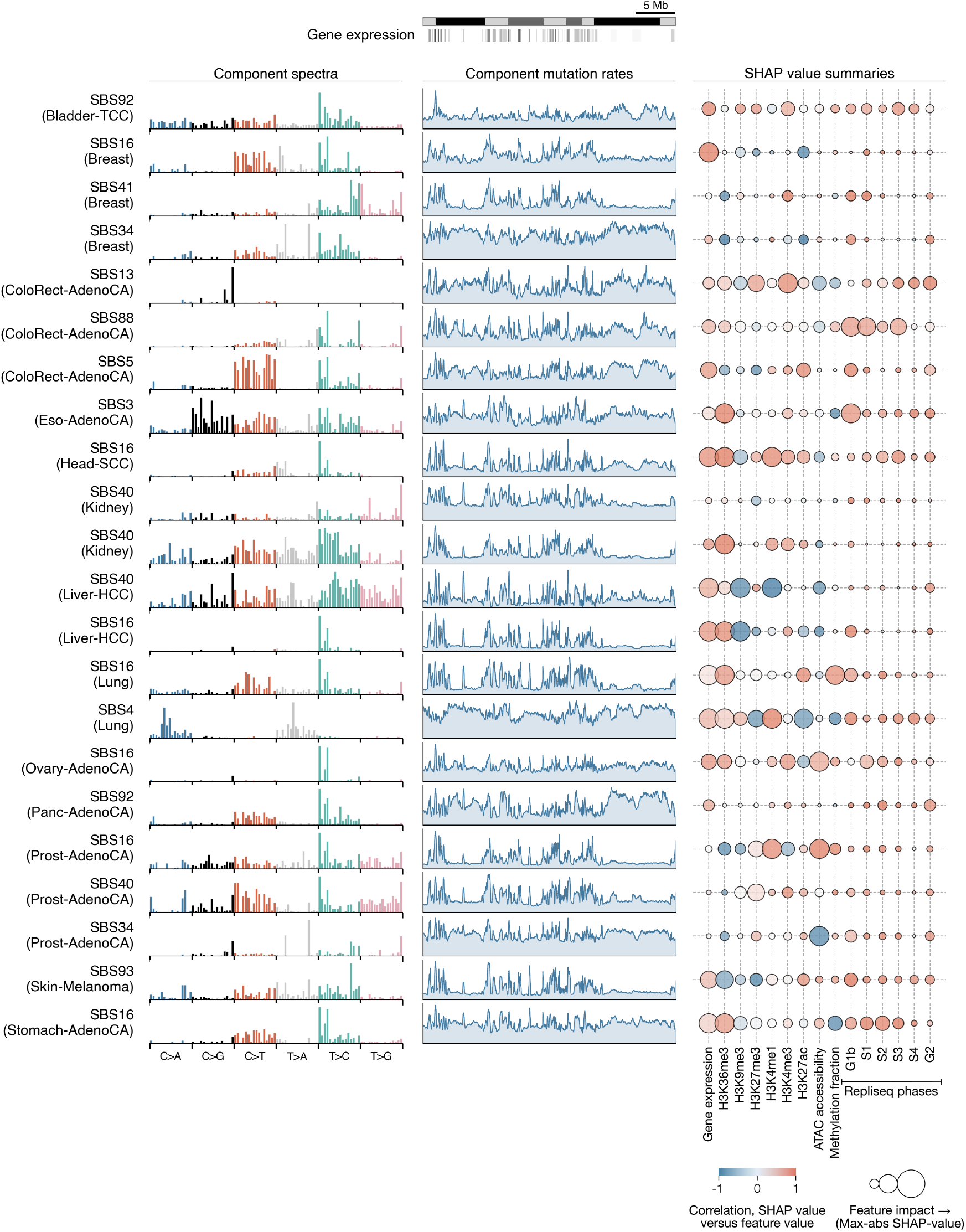
Topo-Tr summary. Mutational spectra, mutation rate profiles, and Shapley value analysis of all signatures in topotype Topo-Tr (transcription).

**Supplementary Figure 16:**
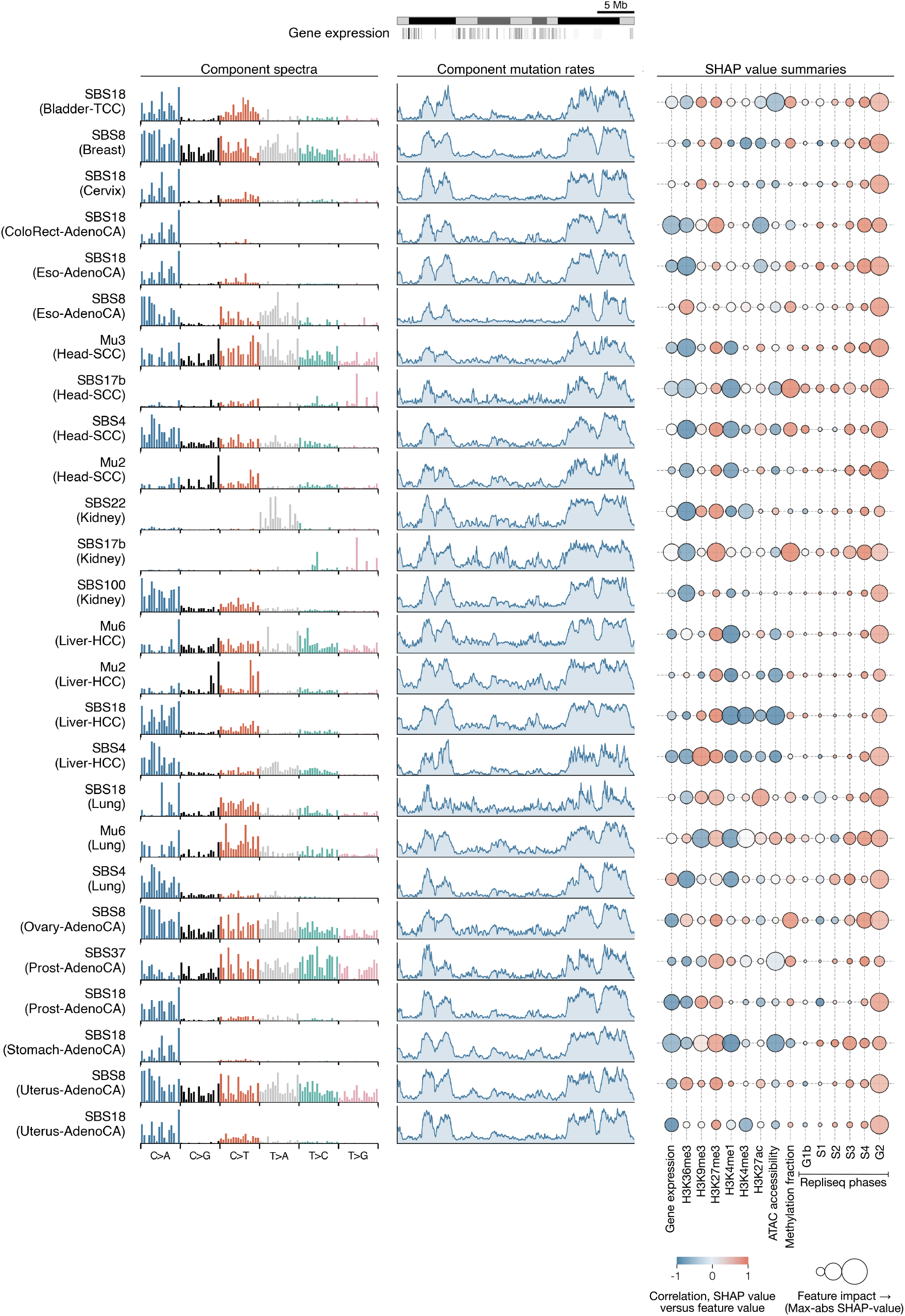
Topo-CanD-I summary. Mutational spectra, mutation rate profiles, and Shapley value analysis of all signatures in topotype Topo-CanD-I (canonical damage I).

**Supplementary Figure 17:**
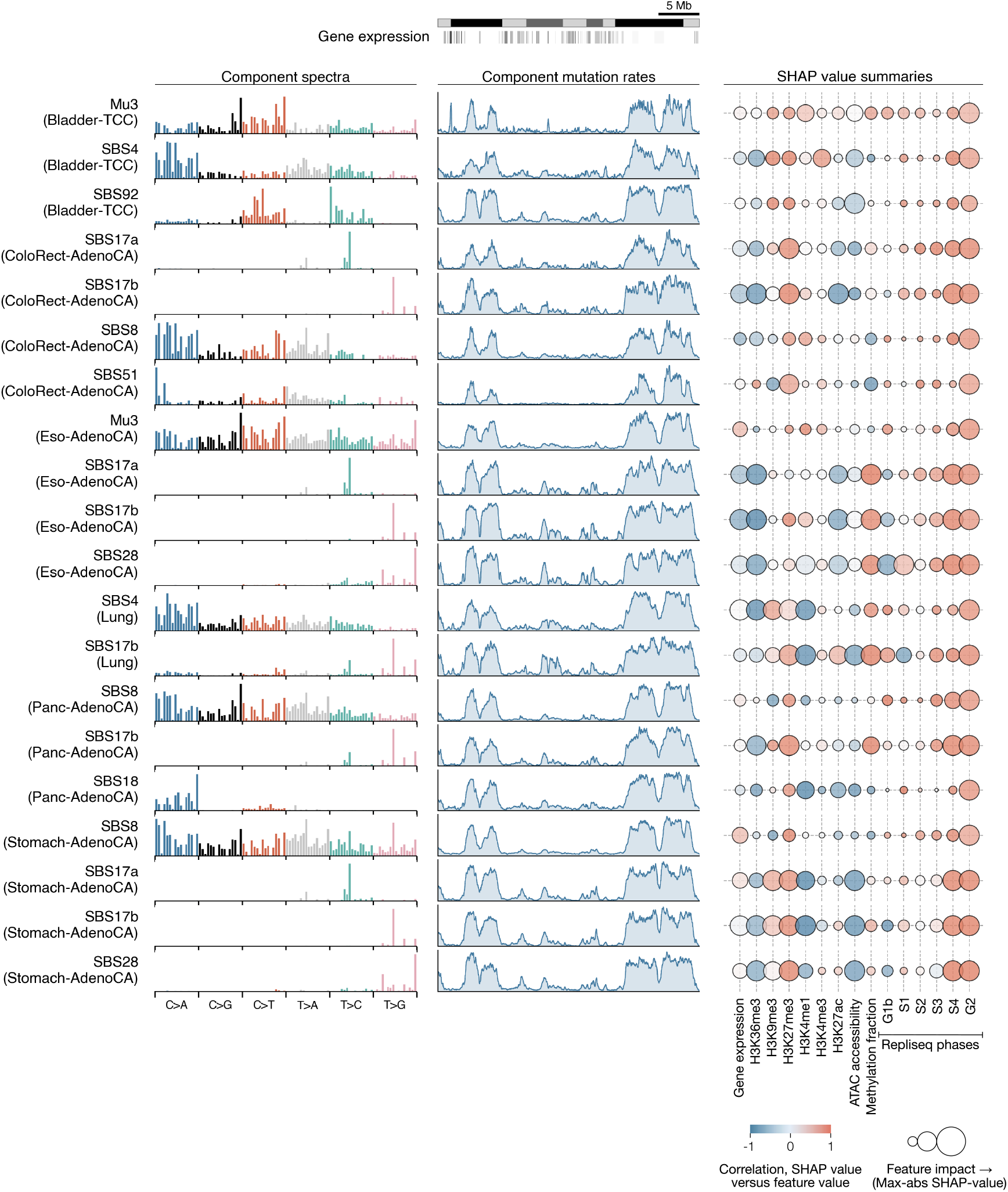
Topo-CanD-II summary. Mutational spectra, mutation rate profiles, and Shapley value analysis of all signatures in topotype Topo-CanD-II (canonical damage II).

**Supplementary Figure 18:**
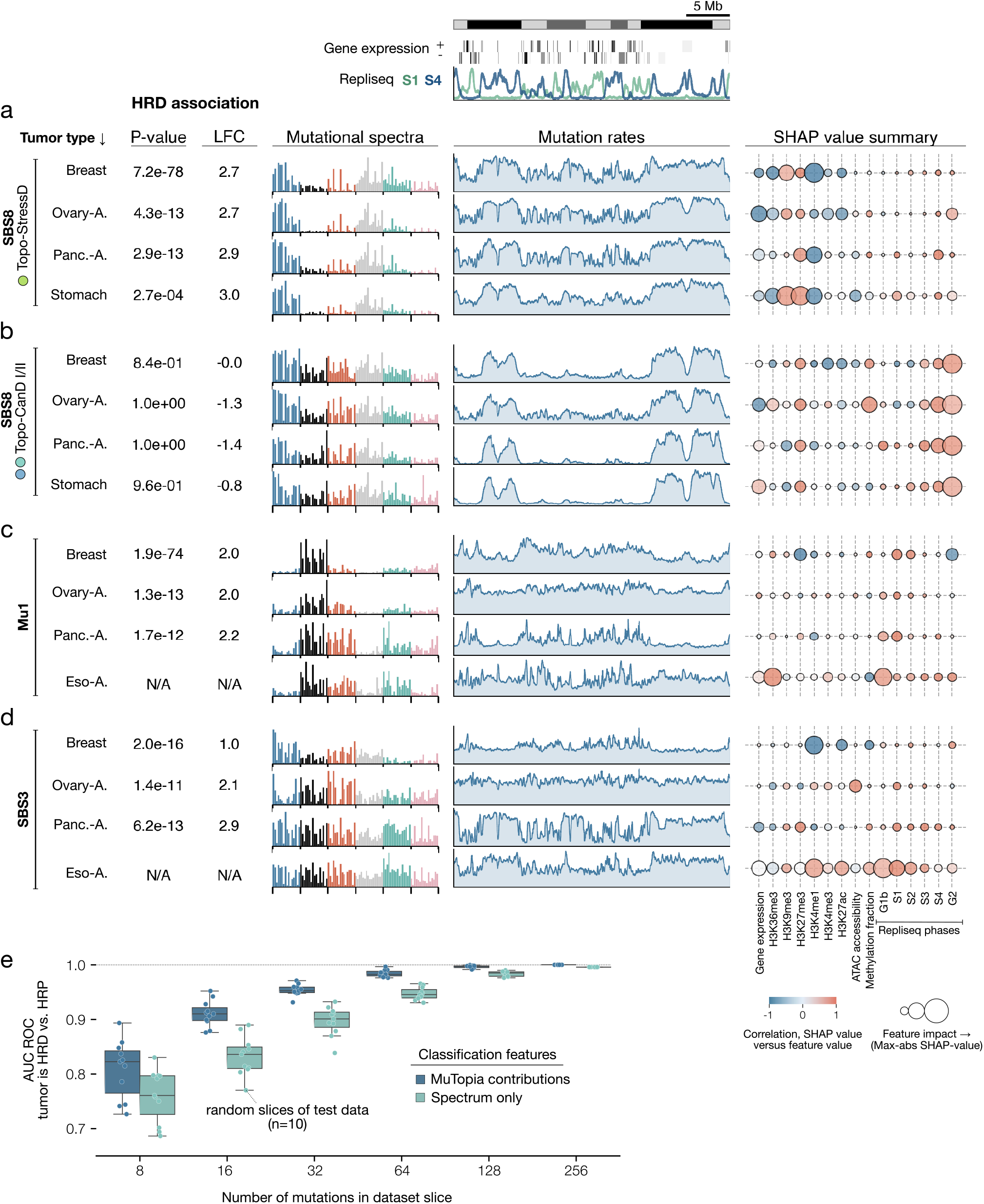
HRD shapes mutational topography. **a)** Mutational spectra, predicted mutation rate profiles, and Shapley value analysis for SBS8 signatures in the Topo-StressD topotype. The p-value and LFC (log2 fold-change) statistics show the enrichment of each process in HRD versus HRP samples, as predicted by CHORD. (Mann–Whitney U-test, Benjamini–Hochberg correction.) **b)** Same as in (a), but for SBS8 signatures in the Topo-CanD-I/II topotypes in the same tumor types. **c)** Same as in (a), but for Mu1 signatures. No esophageal adenocarcinoma samples were called HRD-positive, so these were labeled “N/A”. **d)** Same as in (c), but for SBS3 signatures in the same tumor types. **e)** HRD classification performance of a logistic regression model trained on either the mutational spectra of ovarian adenocarcinoma whole genome samples, or the contributions of each mutational process to the same tumors as inferred by the MuTopia model. We sub-sampled the test-set samples to assess the minimum number of mutations needed for classification using both feature sets.

**Supplementary Figure 19:**
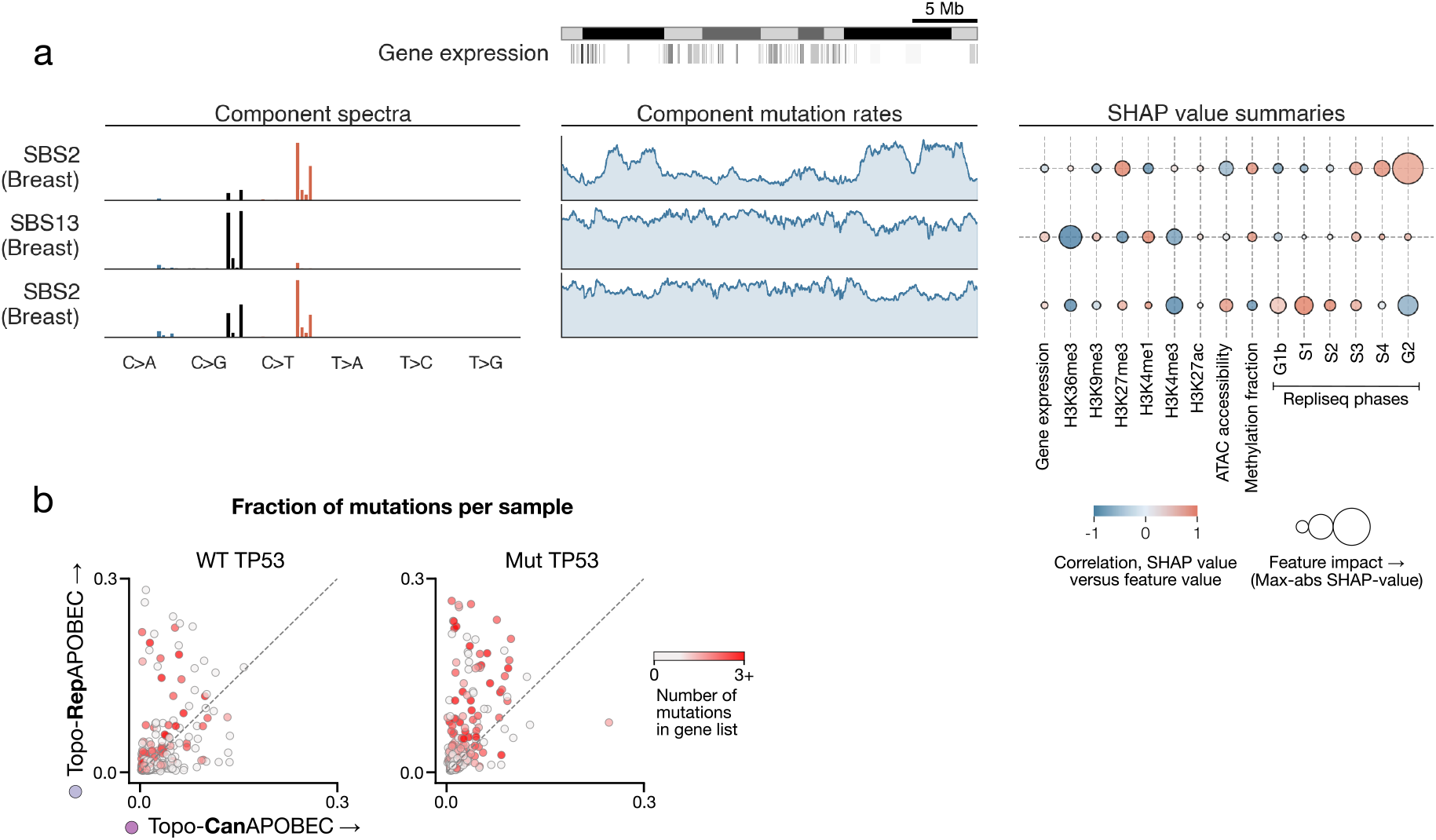
APOBEC exhibits varied genomic mutational profiles. **a)** Mutational spectra, mutation rate profiles, and Shapley value analysis for APOBEC signatures extracted from the breast tumor dataset. **b)** Fraction of mutations per sample attributed to Topo-CanAPOBEC versus Topo-RepAPOBEC, stratified by TP53 mutation status and colored by number of mutations in cancer driver genes.

**Supplementary Figure 20:**
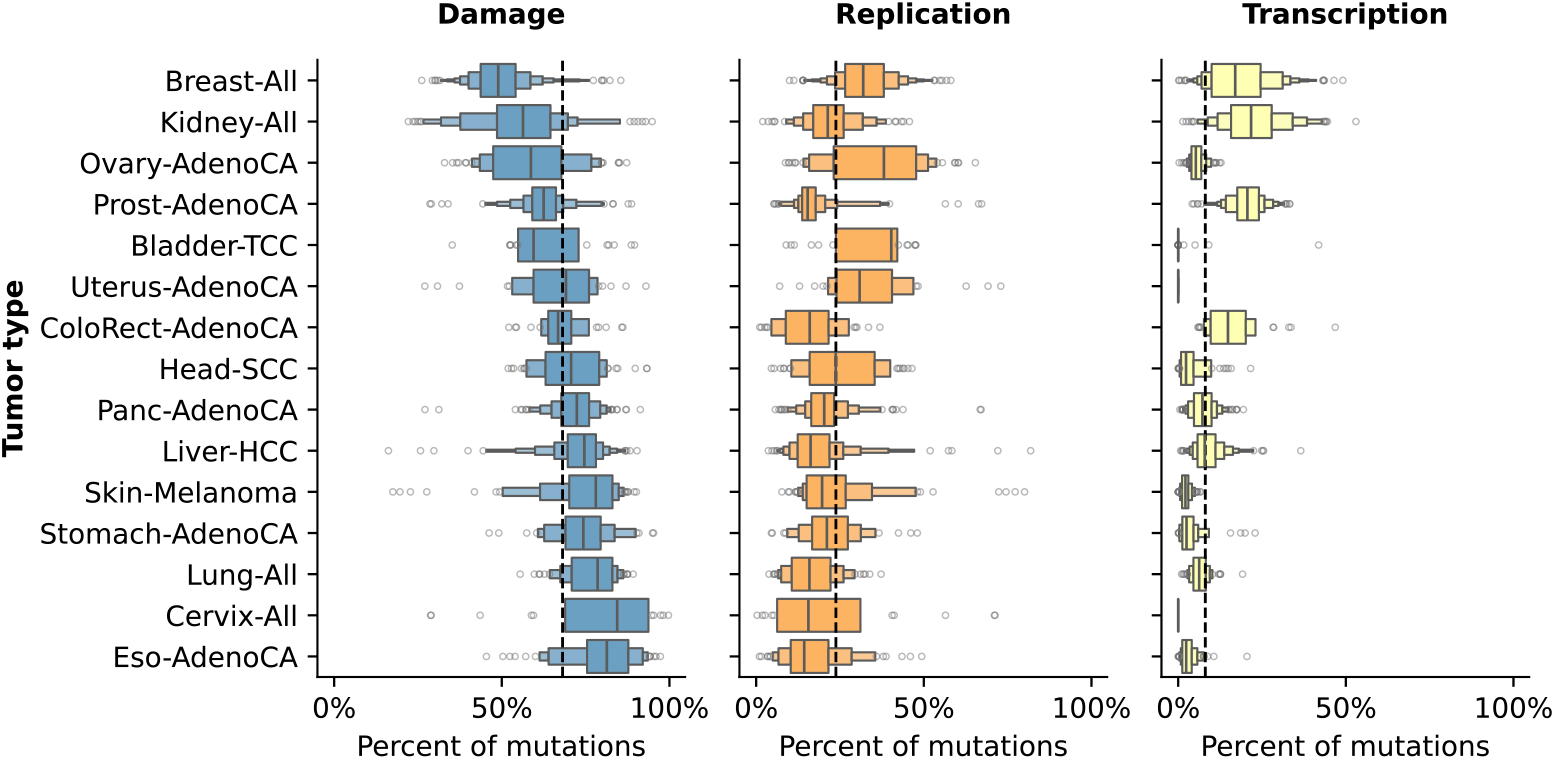
Fractions of damage, replication, and transcription-associated mutagenesis. Relative fraction of mutations from damage topotypes (Topo-CanD-I/II, Topo-CanAPOBEC, Topo-StressD), replication topotypes (Topo-Rep, Topo-RepAPOBEC), and transcription topotypes (Topo-Tr) across samples for each tumor type. The dashed line shows the average across tumor types.

**Supplementary Figure 21:**
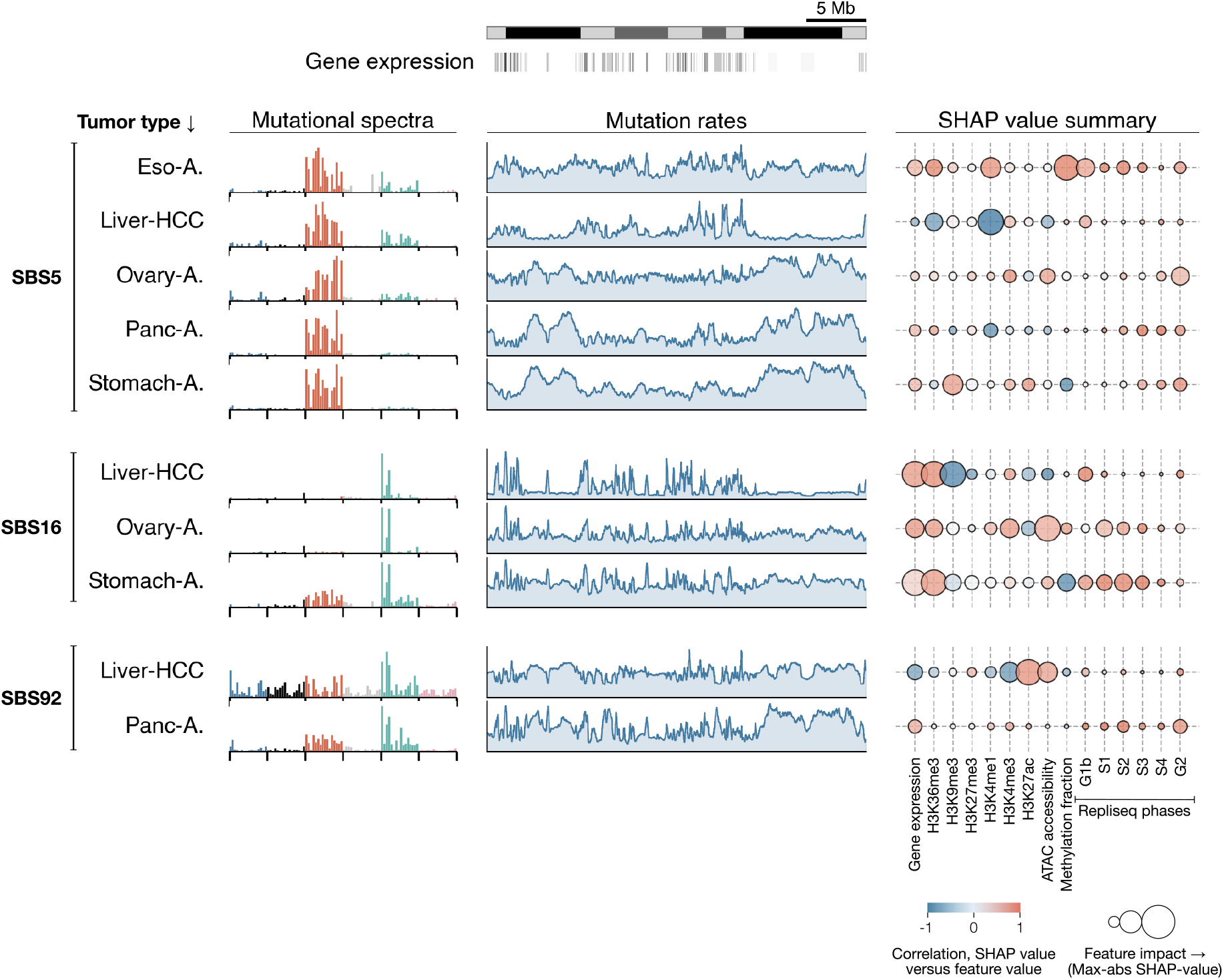
SBS5 as a composite signature of SBS30, SBS16, and SBS92. Mutational spectra, mutation rate profiles, and Shapley value analysis of SBS30-, SBS92-, and SBS16-designated signatures from tumor types which did not contain SBS5.

